# Structural basis of adenylyl cyclase 9 activation

**DOI:** 10.1101/2021.08.11.455989

**Authors:** Chao Qi, Pia Lavriha, Ved Mehta, Basavraj Khanppnavar, Inayathulla Mohammed, Yong Li, Michalis Lazaratos, Jonas V. Schaefer, Birgit Dreier, Andreas Plückthun, Ana-Nicoleta Bondar, Carmen W. Dessauer, Volodymyr M. Korkhov

## Abstract

Adenylyl cyclase 9 (AC9) is a membrane-bound enzyme that converts ATP into cAMP. The enzyme is weakly activated by forskolin, fully activated by the G protein Gαs subunit and is autoinhibited by the AC9 C-terminus. Although our recent structural studies of the AC9-Gαs complex provided the framework for understanding AC9 autoinhibition, the conformational changes that AC9 undergoes in response to activator binding remains poorly understood. Here, we present the cryo-EM structures of AC9 in several distinct states: (i) AC9 bound to a nucleotide inhibitor MANT-GTP, (ii) bound to an artificial activator (DARPin C4) and MANT-GTP, (iii) bound to DARPin C4 and a nucleotide analogue ATPαS, (iv) bound to Gαs and MANT-GTP. The artificial activator DARPin C4 partially activates AC9 by binding at a site that overlaps with the Gαs binding site. Together with the previously observed occluded and forskolin-bound conformations, structural comparisons of AC9 in the four new conformations show that secondary structure rearrangements in the region surrounding the forskolin binding site are essential for AC9 activation.

**One Sentence Summary:** Cryo-EM reveals activator-induced conformational changes in adenylyl cyclase AC9

## Introduction

Adenylyl cyclases (ACs) play a fundamental role in many G protein-coupled receptor (GPCR) mediated signal transduction pathways ^1-3^. Activation of a Gαs-coupled GPCR by an extracellular stimulus, such as a hormone, leads to a cascade of events that include the exchange of GDP bound to the Gαs subunit to GTP, dissociation of the GTP-bound Gαs from the heterotrimeric G protein-GPCR complex, followed by binding of the GTP-bound Gαs to a membrane AC. The interaction with Gαs potentiates the ACs ability to convert a molecule of adenosine 5’-triphosphate (ATP) into cyclic adenosine monophosphate (cAMP) ^1,4^. The produced cAMP is a key second messenger in many living cells, binding to and regulating a number of downstream effector proteins, and thus modulating a plethora of physiological functions ^5^. The nine subtypes of membrane ACs described in mammals, AC1-9, differ in cellular localization, tissue distribution and physiological functions ^5^. For example, the ACs predominantly expressed in the nervous system, AC1 and AC8, are linked to cognitive processes and pain perception ^2^, whereas in heart, AC5, AC6 and AC9 are linked to heart disease ^2,6^. Mutations of several membrane ACs have been linked to genetic diseases, including autosomal deafness 44 (AC1) ^7^, obesity and type 2 diabetes (AC3) ^8^, familial dyskinesia with facial myokymia (AC5) ^9^, or lethal congenital contracture syndrome 8 (AC6) ^10^.

The membrane ACs share a conserved predicted domain arrangement, membrane topology and a high degree of sequence similarity, particularly in the catalytic regions of the protein ^11^. Each membrane AC contains two conserved cytosolic catalytic domains and twelve predicted transmembrane (TM) helices, with cytosolic N- and C-termini of varied lengths and regulatory roles ^1,11^. Insights into the structure and molecular mechanism of the membrane ACs have been gained by the early X-ray crystallographic studies on the chimeric soluble domain of adenylyl cyclase (AC5_c1_/AC2_c2_) in complex with Gαs and forskolin ^12-14^. These studies provided the structural basis for the two metal-ion-catalysis of ATP-cAMP conversion, revealed some of intermediate states of the enzyme, and provided a plausible explanation of enzyme activation by the plant-derived small molecule activator, forskolin ^15^. While forskolin is a non-physiological activator of the membrane ACs, an endogenous molecule that regulates the ACs at the allosteric binding site has not yet been identified. Forskolin and Gαs were required to stabilize the dimer of the isolated AC5_C1_ and AC2_C2_ domains, as both regulators enhance the affinity between C1a and C2a in the soluble system ^12-14^. The mechanism of forskolin-mediated AC activation was ascribed to its ability to “lock” the two domains closer together, likely inducing a conformational change to reorient C1a and C2a with respect to each other to enhance activation ^12-14^. Recently we determined the cryo-EM structure of the full-length bovine AC9 bound to Gαs in an auto-inhibited “occluded” state, together with the structure of a C-terminally truncated AC9 (AC9_1250_), bound to Gαs, forskolin and MANT-GTP, a non-cyclizable AC inhibitor ^1^. These structures provided the first glimpse into the architecture and auto-inhibitory regulation of a full-length membrane AC. Furthermore, the ability of forskolin to activate AC9 has been a controversial subject, with some studies indicating that the enzyme is insensitive to forskolin ^2^. The structure of AC9_1250_-Gαs bound to MANT-GTP and forskolin, combined with biochemical studies, confirmed that AC9 can be activated by forskolin binding to its canonical allosteric site in the presence of Gαs ^1,16^.

AC9 is expressed ubiquitously and plays an important physiological role. A polymorphism of AC9, rs2230739, has been associated with differences in susceptibility to drug treatment in asthma patients’, indicating AC9 as a potential asthma drug target ^2,17^. In the heart, AC9 is involved in the slow delayed rectifier (I_Ks_) current through its interaction with the A-kinase anchoring protein, Yotiao and KCNQ1/KCNE1 channels ^18^. AC9 has also been suggested to have a cardioprotective role in the heart through its interaction with Hsp20 ^6^.

Despite the availability of the high-resolution X-ray structures of the AC5_c1_/AC2_c2_ domains as well as the single particle cryo-EM structures of AC9, the molecular determinants of AC9 regulation remain unclear. We set out to determine the key conformational changes associated with distinct functional states of AC9. The choice of the AC9 as a model was motivated by both its tremendous biomedical significance and relevance to disease, as well as by our ability to probe the structure of the full-length protein. To aid our structural and biochemical analysis we developed an artificial binder of AC9, a designed ankyrin repeat protein (DARPin) ^19^ that binds to AC9 and partially activates it. Here we present four cryo-EM structures: (i) AC9 in complex with MANT-GTP (AC9-M), (ii) AC9 in complex with MANT-GTP and an artificial partial activator, DARPin C4, which is capable of activating the AC9 without inducing the occluded state (AC9-C4-M), (iii) a DARPin C4-bound state with ATPαS instead of MANT-GTP (AC9-C4-A), (iv) AC9_1250_ in complex with Gαs and MANT-GTP, in the absence of forskolin (AC9_1250_-Gαs-M). Analysis of all structures of AC9 available to date reveals the conformational transitions that the protein goes through in response to binding of distinct activating agents (DARPin C4, Gαs, forskolin). The magnitude of the conformational changes correlates with the potency of the activator. Our data suggest that activation of AC9 is intricately linked to the conformation of the allosteric site.

## Results

### Structure of AC9 bound to MANT-GTP

To characterize the conformation of AC9 in the absence of G protein α subunit, we determined the structure of bovine AC9 in complex with MANT-GTP (referred to as AC9-M throughout the text below) using cryo-EM and single particle analysis at 4.9 Å resolution (Supplementary Fig. 1). Our choice of MANT-GTP as a ligand for the active site of AC9 was based on two considerations: (i) we have used it successfully in our previous studies of membrane ACs, (ii) the conformation stabilized by MANT-GTP is very similar to that stabilized by ATP analogues, such as ATPαS, based on X-ray crystallographic studies ^13,20^. Due to the low molecular weight and innate flexibility and pseudo-two-fold symmetry of the full-length AC9, the resolution of the AC9-M reconstruction could not be improved beyond 4.9 Å. Nevertheless, the structure revealed the main features of the protein, including the TM1-12 helices, the helical domain, and the catalytic domain. The positions of these domains could be assigned to the density features unequivocally (Supplementary Fig. 1).

### Artificial partial activator of AC9, DAPRin C4

In order to stabilise the protein, increase its size for cryo-EM analysis and potentially break the pseudo-two-fold symmetry, we selected a panel of designed ankyrin repeat proteins (DARPins) by using ribosome display ^19,21,22^. From a small pool of 21 DARPins that specifically bound to AC9, we identified DARPin C4, capable of activating AC9 alone, but not the AC9-Gαs protein complex (Fig. 1a-b). Analysis of the AC specificity of DARPin C4 revealed that it is highly selective for AC9 (Fig. 1c,e). The affinity of the DAPRin C4 for purified AC9 in detergent micelles, based on AC activity assays, is in the high nanomolar range (Fig. 1d). The cAMP accumulation assays showed a half-maximal effective concentration (EC_50_) for DARPin C4 activation of AC9 of 4.4 nM (8.3 nM for AC9_1250_), with very similar V_max_ values (∼120 nmol/mg/min for both AC9 and bAC9_1250_). These V_max_ values are ∼3-fold higher compared to the basal activity of AC9 (∼37 nmol/mg/min)^1^. In comparison, Gαs activated the full-length AC9 significantly stronger, with the maximal effect seen in the absence of the C2b domain, i.e., in the AC9_1250_ construct (Fig. 1d). Experiments using membrane preparations of insect cells expressing select AC isoforms confirmed the selectivity of DARPin C4 for AC9 (Fig. 1e) and the high affinity of this agent for AC9 under a variety of experimental conditions (i.e., in the presence of Mn^2+^ or Mg^2+^ ions that differentially support the activity of AC9; Supplementary Fig. 2).

**Fig 1:**
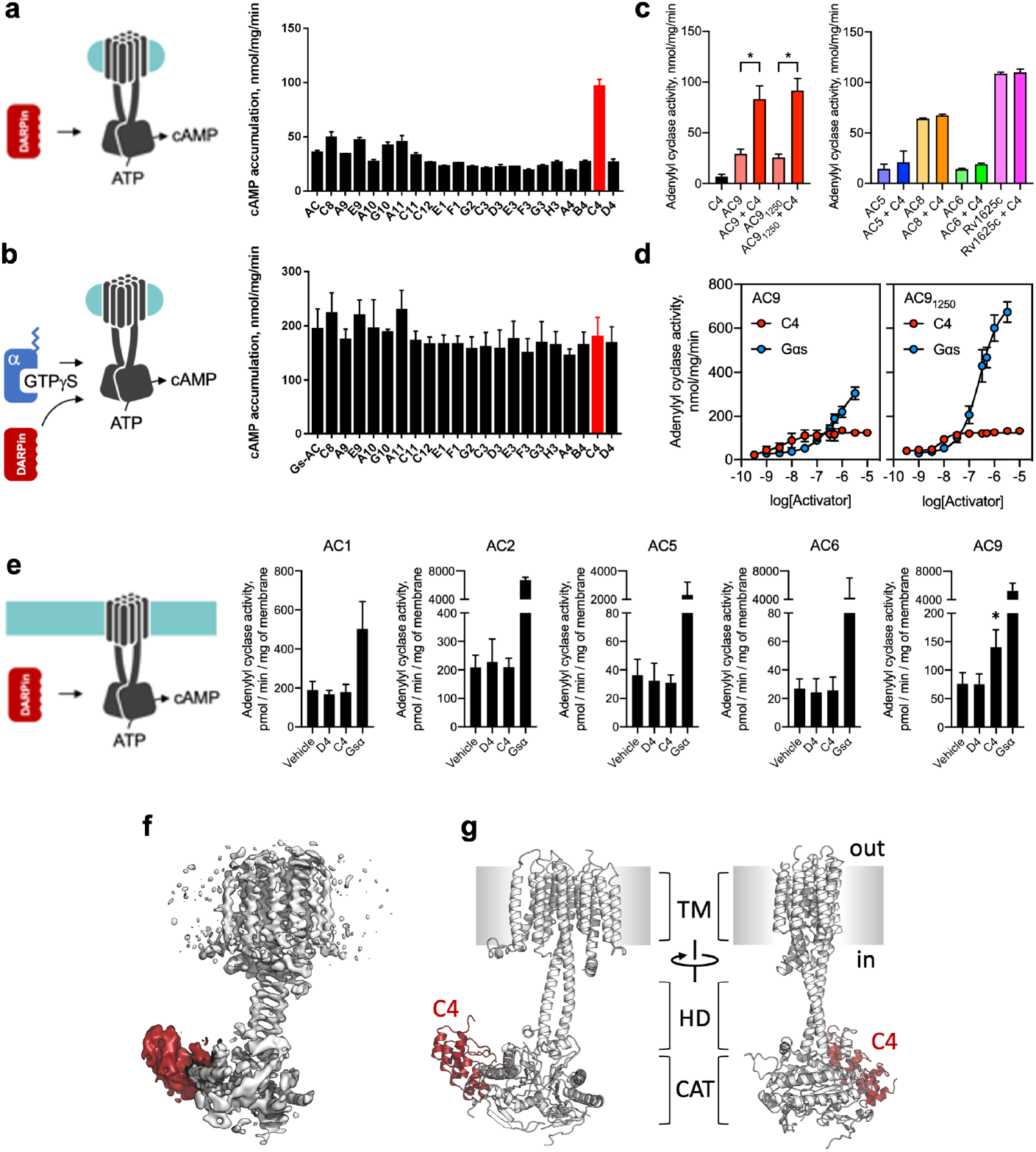
Identification, biochemical properties and structure of DARPin C4 as a partial activator of AC9. **a-b**, AC9 activity screens identify the DARPin C4 as an AC9 activator; the experiment in B was performed in the presence of GTPγS-bound Gαs (n = 2). **c**, Activity of purified, detergent-solubilized ACs (as in a-b) demonstrate selectivity of DARPin C4 for AC9 (n = 3; P < 0.005 indicated by an asterisk). **d**, Dose-response activity curves of the DARPin C4 show that it has a high apparent affinity for AC9 (left) and for AC9_1250_ (right; n = 3); unlike Gαs, DARPin C4 acts as a partial activator of AC9 regardless of the presence of the C2b domain. **e**, AC activity assays performed using membrane preparations of Sf9 cells expressing different ACs confirm that activation by DARPin C4 is selective for AC9. **f**, Cryo-EM map of the DARPin C4-bound AC9 (white map corresponds to AC9, red – DARPin C4. **g**, Model of the AC9-C4 complex; key elements of the structure, including the transmembrane domain bundle (TM), the helical domain (HD), the catalytic domain (CAT) and the DARPin C4 (C4, red) are indicated.

### In vivo effects of DARPin C4

To determine whether DARPin C4 can interact with AC9 in a cellular context, we used FRET microscopy. Analysis of FRET between the YFP-tagged AC9 and CFP-tagged DARPin C4 confirmed the interaction between these proteins (Supplementary Fig. 3). Furthermore, expression of DARPin C4 in HEK293F cells led to increased cAMP accumulation when co-expressed with AC9, compared to control cells (Supplementary Fig. 3). These proof-of-concept experiments established unequivocally that DARPin C4 interacts with and activates AC9 in cells, and thus can be used in cell-based applications.

### Structure of AC9 bound to DARPin C4

To elucidate the structural basis of AC9 activation by DARPin C4, we copurified these two proteins using size exclusion chromatography (SEC) (Supplementary Fig. 4) and subjected the AC9-C4 complex to cryo-EM imaging. Using single particle analysis of the complex we obtained the 3D reconstruction of AC9-C4-M (AC9-C4 complex in the presence of 0.5 mM MANT-GTP) at a resolution of 4.2 Å (Fig. 1f-g, Supplementary Fig. 5-6). The structure revealed all the previously observed features of the AC9 (Fig. 1g), including the 12-TM domain region (TM), the helical domain (HD) and the catalytic domain (CAT). The DARPin C4 binds to the C2a domain of AC9 at a site that partially overlaps with the Gαs-binding site (Fig. 1f, g, Fig. 2a-g). The variable region of DARPin C4 inserts into the groove formed by the α2’ and α3’ helices of the C2a domain of AC9 (Fig. 2a,f), similarly to the way the switch II region of Gαs binds to and activates the enzyme ^1^.

**Fig 2:**
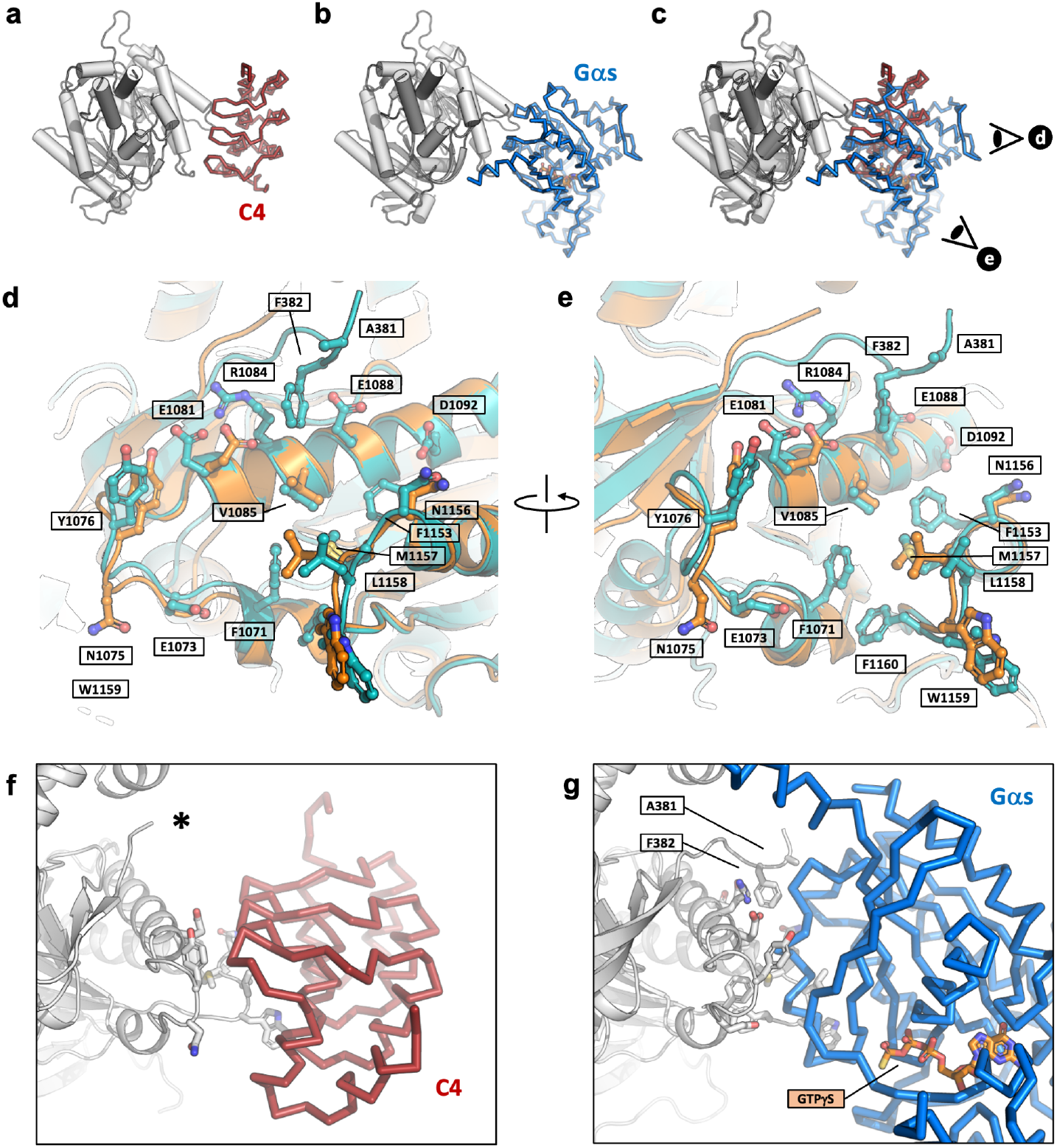
DARPin C4-binding site within the catalytic domain of AC9. **a-c**, Views of the DARPin-C4- (a) and Gαs-bound AC9 (b). **c**. The binding sites of the two interaction partners of AC9 overlap. The symbols in C indicate the points of view in d and e. **d-e**, The views of the activator binding sites in the DARPin C4- (orange) or Gαs-bound (blue) AC9 complexes. The side-chain of all residues within 4 Å of the interaction partner are shown as sticks. **f-g**, The views of the AC9-activator interface, revealing the absence (f, indicated by an asterisk) and the presence (g) of interaction between AC9 and DARPin C4 and Gαs, respectively, involving the C1a domain loop region (residues A381 and F382 in proximity with G protein are indicated).

### Determinants of AC9 activation by DARPin C4

Both activator proteins, Gαs and DARP C4 bind to a very similar region of the C2a domain (Fig. 2d-e), yet their effects on the functional state of the protein are drastically different. There is one major structural difference between the structure of AC9-C4-M and that of AC9-Gαs determined previously ^1^: DARPin C4 does not appear to induce the occluded state, in which the C2b domain of AC9 displaces the nucleotide and forskolin from their binding sites at the C1a/C2a domain interface. In contrast, Gαs induced the I1263-P1275 region of the C2b domain to wedge itself into the substrate and allosteric binding sites of AC9, presumably as part of an auto-inhibitory mechanism that prevents over-production of cAMP ^1,23^.

The remarkable difference between the ability of the DARPin C4 and Gαs protein to activate AC9 and to induce the auto-inhibitory occluded conformation shows that there is a fundamental difference between the molecular interactions of these two proteins. Comparisons of the binding interfaces between AC9 and its regulators, Gαs and DARPin C4, provide clues to the mechanism of AC9 activation by the DARPin C4. A subset of shared residues between the G protein and the DARPin C4-binding sites in the C2a domain, unique to AC9, is likely responsible for the ability of the DARPin C4 to specifically recognize this enzyme (Supplementary Fig. 7a-c). A loop in the C1a domain corresponding to residues I380-P384, appears to be flexible and could not be resolved in the AC9-C4-M map (Fig. 2f). However, this loop does interact with the G protein, evident from the AC9-Gαs complex (Fig. 2g) and the available AC5_c1_/AC2_c2_ crystal structures. This extensive interaction with the regions in both C1a and C2a domain of AC9 makes Gαs a more potent activator, compared to DARPin C4.

The structure of AC9-C4-M revealed density in the active site consistent with the presence of MANT-GTP, as well as a lack of any density in the allosteric site (Fig. 3a). Although the AC9-C4 structure provides clues about elements of AC9 participating in selective binding of DARPin C4, it is unclear why the occluded state of AC9 has only been observed in the presence of Gαs. Moreover, we could only obtain an activated state of the protein bound to Gαs upon removal of the C-terminal C2b domain. In contrast, in the case of the DARPin C4-bound state, the full-length AC9 could be captured in a nucleotide-bound state. It is apparent that there is an intricate link between the Gαs protein and the largely unstructured C-terminus of AC9, which will require further examination.

**Fig 3:**
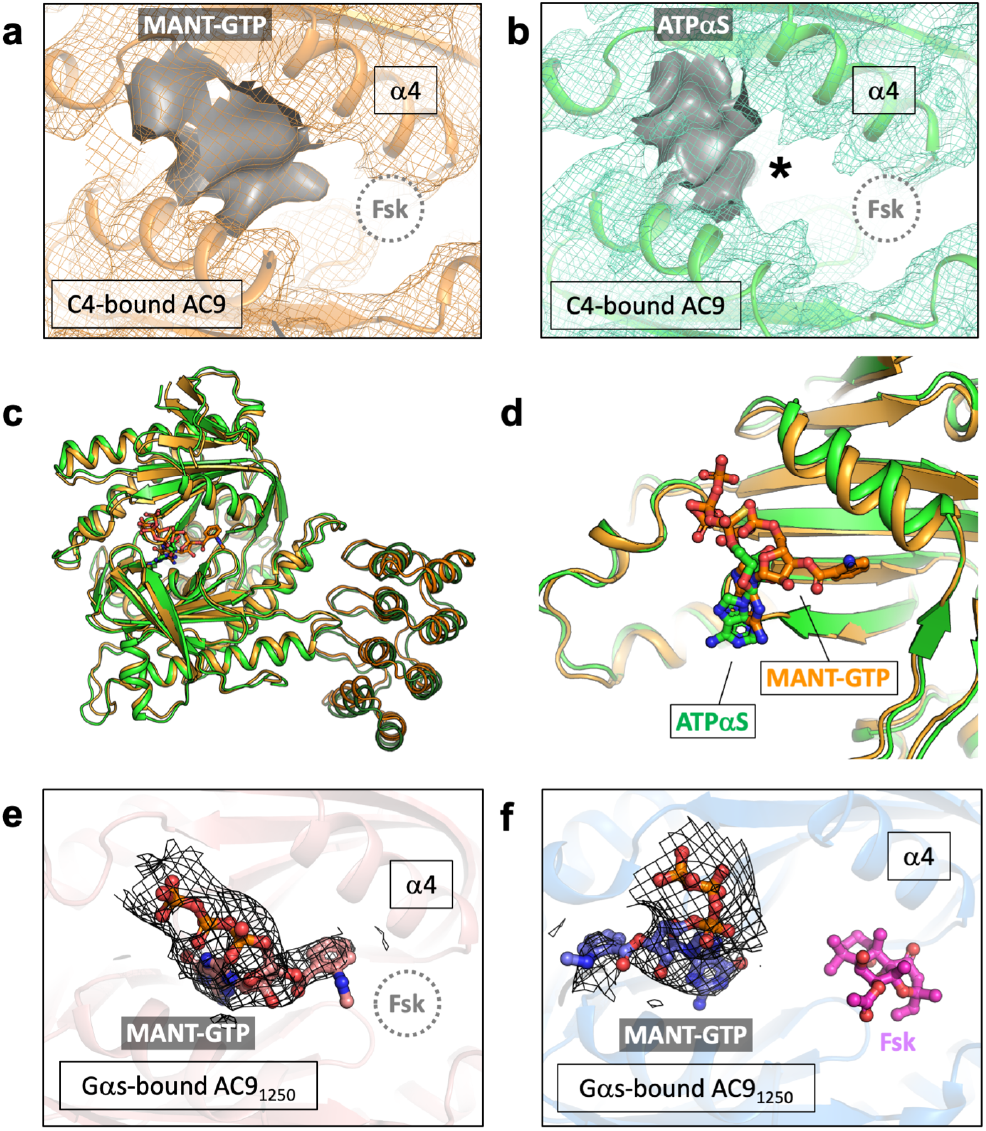
Experimentally observed density in the nucleotide-binding site of AC9. **a**, The portion of the density corresponding to the bound nucleotide (MANT-GTP) in the map AC9-C4 is shown as grey surface; the position of the forskolin binding site is indicated with a dashed circle.**b**, A similar view of the AC9-C4 in complex with ATPαS. The asterisk indicates the absence of extended density. **c**, Alignment of the soluble domains of AC9 in a complex with C4 and MANT-GTP (orange) or ATPαS (green) reveal a low RMSD of 1Å. **d**, Although the conformation of the catalytic domain is very similar, the pose of the nucleotide is different based on the density (as in a-b), with MANT-group of MANT-GTP pointing towards the forskolin site. **e**, The density map corresponding to the bound MANT-GTP molecule in the AC9_1250_-Gαs-M structure (forskolin-free). The density is similar to that in a. **f**, The density corresponding to MANT-GTP in the AC9_1250_-Gαs-MF structure is shown as black mesh. The molecule of forskolin (Fsk) is shown as magenta sticks.

### AC9-bound nucleotide conformation in the absence of forskolin

Although the cryo-EM maps of AC9-M and AC9-C4-M featured the density elements corresponding to a bound MANT-GTP molecule (Fig.3a), the location of the nucleotide density in each of the maps was distinct from that observed in the previously determined AC9_1250_-Gαs structure in the presence of MANT-GTP and forskolin (AC9_1250_-Gαs-MF) (Fig. 3f) ^1^. The nucleotide density stretched out towards to the forskolin binding site, suggesting that the molecule may have been captured in a new orientation within the active site (Fig. 3a). The local resolution of this region in both density maps (AC9-M and AC9-C4-M) was not sufficiently high to unambiguously define the pose of MANT-GTP.

To further structurally characterize the nucleotide binding site of AC9, we determined the structure of AC9_1250_-Gαs-M (the truncated AC9_1250_ in complex with Gαs and MANT-GTP). Focused refinement of the 3D reconstruction improved the resolution of the map corresponding to the soluble part of the complex to 3.8 Å (Supplementary Fig. 8, 9). As in the case of AC9-M and AC9-C4-M, the AC9_1250_-Gαs-M map featured the density of the nucleotide extending towards the allosteric site (Fig. 3a,e). The previously determined nucleotide pose “M1” corresponds to the structures of AC9_1250_-Gαs-MF ^1^ (Fig. 3f), the AC5_c1_/AC2_c2_-Gαs ^20^ and the *M. intracellulare* Cya ^24^ (Supplementary Fig. 10a-b). The AC catalytic site is known to be capable of accommodating the nucleotides in non-canonical orientations, evident from two distinct poses of the MANT-GTP in the mycobacterial Cya ^24^ (Supplementary Fig. 10b), the structure of sAC bound to ApCpp and an inhibitor LRE1 ^25^ (Supplementary Fig. 10d) or the structure of *M. tuberculosis* MA1120 bound to Ca^2+^ and ATP ^26^ (Supplementary Fig. 10e). The density occupying the nucleotide binding site observed in AC9-C4-M and AC9_1250_-Gαs-M maps is consistent with a previously undescribed non-canonical pose of MANT-GTP, featuring the MANT group stretching out towards the forskolin binding site.

This unexpected observation prompted us to further investigate nucleotide binding to AC9. Although MANT-GTP is a nucleotide-derived inhibitor, its structure is quite distinct from the natural AC substrate ATP. To better understand the observed extended density protruding from the nucleotide binding site, we determined the structure of AC9 in a complex with DARPin C4 and an ATP analogue ATPαS (AC9-C4-A) at 4.3 Å resolution (Supplementary Fig. 11, 12). While ATPαS is also an AC inhibitor, it lacks the MANT-group and is likely a better representative of the nucleotide pose in the ATP-bound state of AC9. The structure of AC9-C4-A is very similar to AC9-C4-M, with the full model and soluble domain RMSD of ∼2 Å and ∼1 Å, respectively. The absence of the stretched density observed in all of the forskolin-free MANT-GTP-bound structures (Fig.3b) indicates that the stretched density in AC9-C4-M and AC9_1250_-Gαs-M map corresponds to the MANT group. We propose that MANT-GTP in the absence of forskolin adopts a stable MANT-GTP pose “M2”, which is distinct from the M1 pose and is characterized by the MANT-group extending towards the forskolin site.

To determine whether AC9 may have any preference for a specific nucleotide pose and to ascertain that the M2 state may represent a stable nucleotide-bound conformation, we performed molecular dynamics (MD) simulations using the cytosolic portion of AC9-Gαs model, substituting the MANT-GTP molecules in M1 and M2 poses with the molecules of ATP placed in the corresponding poses ATP1 and ATP2 (Supplementary Fig. 13a-d). During the MD simulations molecules placed at ATP2 remained bound at their starting locations (the RMSDs of the nucleotide atoms were around 2 Å; Supplementary Fig. 13d; Movie 2). Molecules at ATP1 shifted from their initial location (Supplementary Fig. 13c; Movie 1).

Taken together, the simulations performed suggest that location M1, albeit favourable for MANT-GTP, is somewhat unfavourable as an ATP-binding site. It is likely that MANT-GTP, an AC inhibitor, is stabilized in the corresponding M1 conformation through additional π-π stacking interactions of the MANT group (e.g., with the W1188 residue in AC9).

### AC9 conformations induced by distinct activators

The availability of AC9 structures captured in distinct conformations elicited by different activating agents (Gαs protein, DARPin C4, forskolin) allows us to determine the elements of AC9 structure that respond to the interaction with these activators (Fig. 4, 5). We aligned AC9-Gαs, AC9-C4-M, AC9_1250_-Gαs-M, AC9_1250_- Gαs-MF to AC9-M, using the C2a domain as an alignment template (Fig. 4, Movie 3). Binding of DARPin C4 to the C2a domain of AC9 leads to a small rearrangement of the C1a domain (RMSD ∼1.2 Å), particularly prominent at the flexible “claw” region (β6α5β7β8) of C1a domain (Fig. 4b). Close to the “claw” region, the helix α4 in AC9-C4-M structure also moves outward (RMSD ∼1.7 Å) compared to AC9. A similar conformational change is also observed in the AC9-C4-A structure, indicating that the overall domain rearrangement is primarily driven by the DARPin C4 rather than by the ligand occupying the active site of the cyclase (Supplementary Fig. 14). The helix α4 lines the forskolin binding interface and has direct interactions with forskolin. In the AC9-Gαs structures, this α4 helix is displaced further (Fig. 4c-d), with RMSD values of 3.15 Å in AC9_1250_-Gαs-M and 5.7 Å in AC9_1250_-Gαs-MF. Likewise, the helix α4 in the occluded state of AC9-Gαs shows an RMSD of 5.0 Å, compared to the AC9-M structure (Fig. 4a). This comparison establishes a clear pattern of conformational changes that correlate with the activation state of the AC9, where helix α4 appears to move out and create an opening in the allosteric site of AC9 concomitantly with the increase in the catalytic activity of the enzyme (Fig. 4, 5). This opening is relatively small in the partially activated state (AC9-C4), and becomes substantial in the forskolin-free, G protein-bound state of AC9 (AC9_1250_-Gαs-M). It is likely to be further stabilized in the presence of forskolin (AC9_1250_-Gαs-MF).

**Fig 4:**
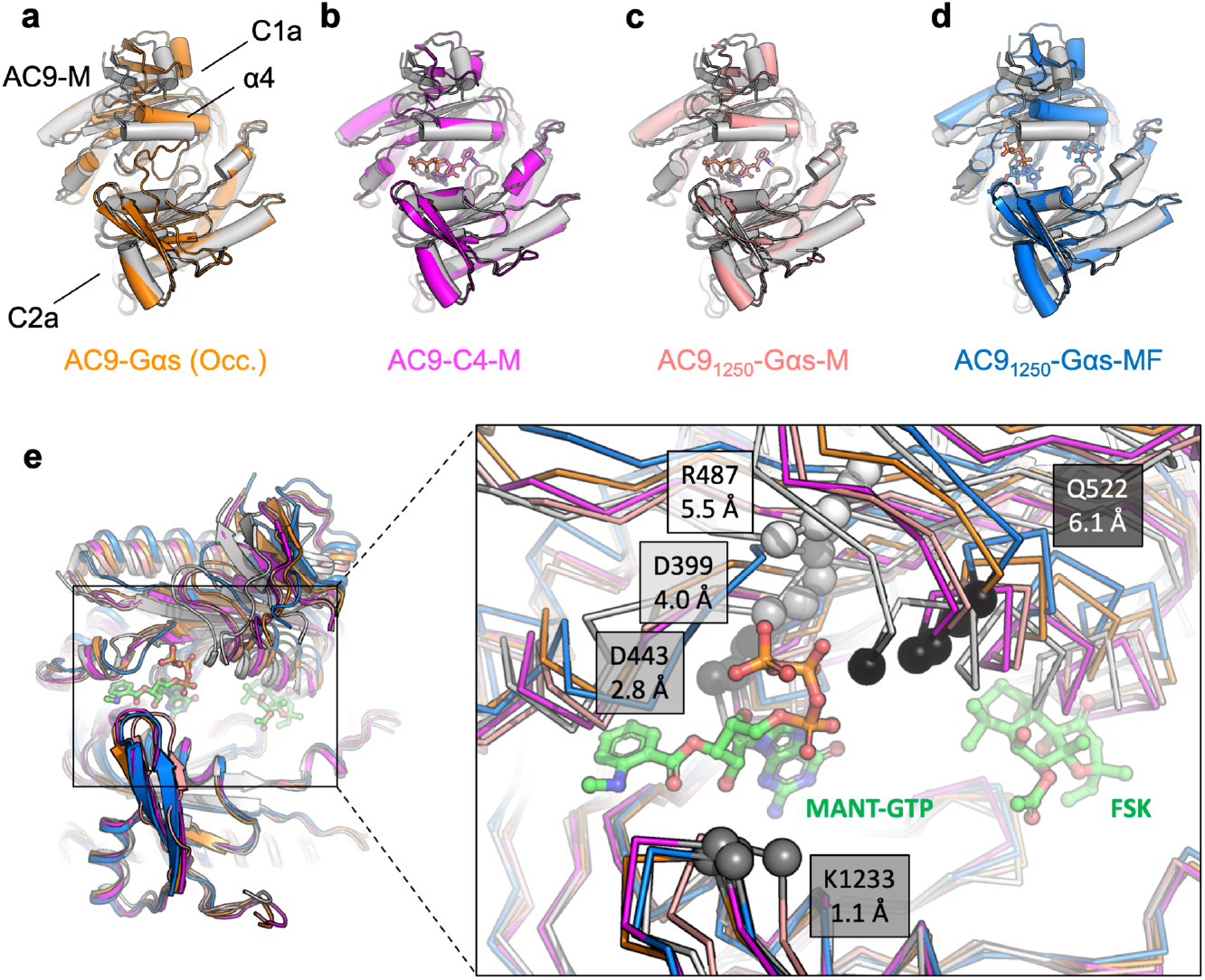
Structural transitions corresponding to the distinct activator-bound states of AC9. **a**, The AC9-M structure (white) was structurally aligned to AC9-Gαs (orange) using the C2a domain as an anchor. The helix α4 is indicated in the panel. **b-d**, Same as A, with the AC9 bound to DARPin C4 (B; magenta), AC9_1250_-Gαs-M (C; pink) and AC9_1250_-Gαs-MF (D; blue). **e**, Comparison of the five available structures reveals relative displacement of the active site residues, D399, D443, R487 and K1233, as well as the Q522 residue in the helix α4 adjacent to the forskolin site. The rearrangement of the active site residues proceeds in a manner that correlates with the activation state of the protein (additionally illustrated by the morphs in Movie S3). *Inset*: The values of Cα atom displacement from the AC9-M (white ribbon) to AC9_1250_-Gαs-MF (fully activated state, blue ribbon) are indicated in grey boxes (Å).

**Fig 5:**
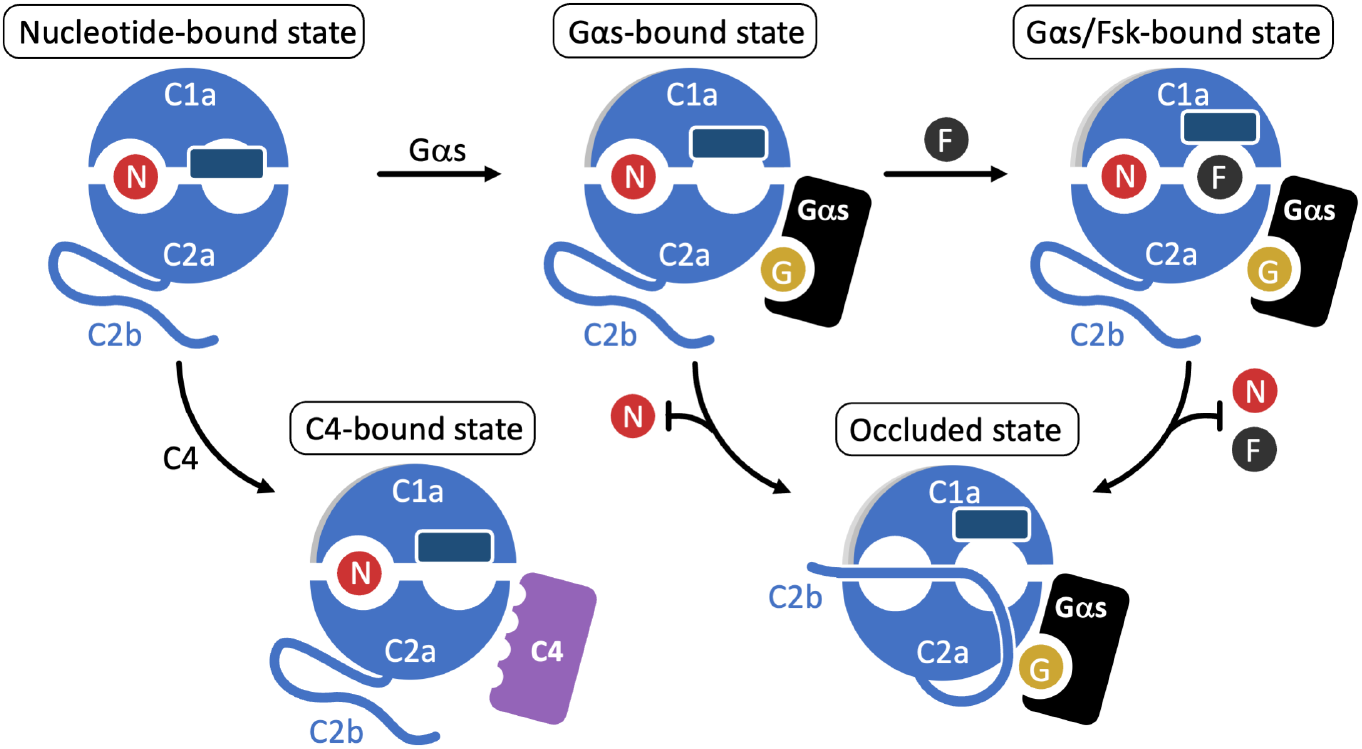
Conformations observed in the available AC9 structures. Based on the available structures, the G protein-free state (nucleotide-bound state, corresponding to AC9-M) can transition to a partially active state upon Gαs binding (Gαs-bound state); subsequent binding of forskolin (Fsk) fully activates AC9 (Gαs/Fsk-bound state). The G protein-activated states can be inhibited by the C2b domain of AC9 with formation of the occluded state. A partially active state can be formed upon DARPin C4 binding. This conformation of AC9 corresponds to a partially activated state, but it is a state that does not favour the formation of the stable occluded conformation, based on the structural evidence. Circles labelled “N”, “G” and “F” correspond to ATP or an ATP substitute (MANT-GTP or ATPαS), GTP and forskolin, respectively. The dark-blue rectangle corresponds to the helix α4, adjacent to the forskolin binding site.

## Discussion

DARPins are established as powerful agents for stabilization of macromolecules for structural and biochemical studies ^19,27^. Although our primary motivation to generate DARPins that bind AC9 was to facilitate AC9 structure determination, the discovery of DARPin C4 as a partial AC9 activator has provided us with a unique opportunity to probe the structure and function of AC9. This reagent has proven to be an excellent tool for studying the molecular determinants of AC9 activation and auto-inhibition. The inability of DARPin C4 to induce the occluded state of AC9 points to a deep link between the specific activation of AC9 by the G protein αs subunit and the built-in auto-regulatory mechanism for controlling the cAMP generation by this membrane enzyme. The auto-regulatory mechanism may be phosphorylation-dependent, based on the existing biochemical evidence ^23^. It is possible that the difference in the interaction interface between DARPin C4 and Gαs accounts for the ability of the G protein to trigger occlusion of the active and allosteric site by the AC9 C-terminus. It is also possible that the disordered regions of AC9, invisible in our 3D reconstruction, play a role in the enzyme activation and our results hint at a yet undescribed interplay between the unstructured loop regions (e.g., N- and C-terminus, C1b region) of the membrane ACs and their regulators, such as the Gαs protein.

The prominent conformational changes involving the helix α4 of AC9 draw parallels to the mechanism of soluble AC (sAC) activation ^28^. In sAC, bicarbonate binding to the allosteric site (approximately corresponding to the forskolin site in the membrane ACs) induces the conformational change of the side chain of R176 residue located on α4. The rearrangement of R176 releases the side chain of D99, acting as a switch that enables the formation of the catalytic cation-binding site (Supplementary Fig. 15). The helix α4 in sAC and its equivalent in AC9 appear to be linked the enzymatic activation mechanism. Activation of sAC is controlled by minute side-chain rearrangement in R176 in response to its activator (bicarbonate). In contrast, AC9 reacts to its activators with whole domain rearrangements, with displacement of helix α4 corresponding to the potency of the activator. The relatively low resolution of our cryo-EM structures could not allow us to determine the side chain orientations in α4 unambiguously, and the precise residues in control of the activation mechanism will require detailed future analysis at higher resolution.

The structures of AC9 available to date have several recognized limitations: (i) The majority of the structures were obtained at a relatively low resolution (∼4Å); (ii) Many of the structures could only be determined in the presence of MANT-GTP, a nucleotide-derived AC inhibitor. The similarity of the MANT-GTP and ATPαS-bound state of AC9-C4 complex indicates that comparisons of distinct MANT-GTP-bound states provide a valid interpretation of the conformational transitions induced by different AC9 activators in the presence of MANT-GTP as a nucleotide substitute, at least at the resolution of our 3D reconstructions. Currently we are severely limited in our ability to probe the structure and function of membrane ACs due to their modular organization and high degree of flexibility. Thus, the use of minimally modified ATP analogues that closely match the structure of ATP as well as development of novel strategies for AC9 stabilization for structural studies will be required to provide new insights into the mechanism of AC9 activation at atomic resolution.

Our experiments establish DAPRin C4 as a highly valuable molecular tool that can be used for *in vitro* biochemistry or in cell-based assays. This reagent can not only be used to obtain unique insights into AC9 activation, but it may provide new opportunities for probing distinct aspects of the cAMP signaling pathway by directly modulating the AC9 activity. Although we limited our experiments to the proof-of-concept assays in HEK293 cells, it is possible that DARPin C4 can be used in a wide variety of cellular contexts, in diverse tissues and at the whole-organism level. The properties of DARPin C4 may need to be modified for optimal performance in specific application. For example, targeting DAPRin C4 to the membrane by a lipid- or peptide-based anchor may dramatically increase its potency *in vivo*. Different methods of DARPin C4 delivery to the cytosol should also be explored. Nevertheless, as DARPins are highly amenable to protein engineering, DARPin C4 may be a good start for generating similar reagents with new AC selectivity profiles, which may find use in a wide range of applications.

## Acknowledgements

We thank the PSI EM Facility (Emiliya Poghossian and Elisabeth Müller-Gubler) and the Electron Microscopy Core Facility at EMBL Heidelberg. We thank Felix Weis (EMBL Heidelberg) for the support in high resolution cryo-EM data collection. We thank Spencer Bliven and Marc Caubet Serrabou (PSI) for high performance computing support. We thank Thomas Reinberg, Sven Furler, Cristian Thom and Joana Marinho for performing the ribosome display selection and screening at the HT-BSF unit at the University of Zurich. This study was supported by iNEXT, the Swiss National Science Foundation (VMK, SNF Professorship, 150665 & 176992), Horten Foundation grant (VMK), and a National Institutes of Health Grant RO1 GM060419 (CWD). The cryo-EM density maps have been deposited in the Electron Microscopy Data Bank, with accession numbers EMD-13330, EMD-13331, EMD-13334, EMD-13335, EMD-13336, EMD-13337 and EMD-13338. The coordinates have been deposited in the Protein Data Bank, with entry codes 7PD4, 7PD8, 7PDD, 7PDE, 7PDF, 7PDG and 7PDH.

