## Supplementary information for "Structural basis of adenylyl cyclase 9 activation"

### Methods

#### DARPin generation

To generate DARPin binders against AC9 and the biotinylated C2a domain alternating on either MyOne T1 streptavidin-coated beads (Pierce) or Sera-Mag neutravidin-coated beads (GE) depending on the particular selection round. Ribosome display selections were performed essentially as described (*1*), using a semi-automatic KingFisher Flex MTP96 well instrument. The DARPin library included a mix of N3C-DARPins with randomized and non-randomized N- and C-caps (*2, 3*), respectively, and the successively enriched pools were cloned as intermediates into a ribosome display specific vector (*4*). Selections were performed over four rounds with decreasing target concentration and increasing washing steps to enrich for binders with high affinities. The final enriched pool was cloned as fusion into a bacterial pQE30 derivative vector with a N-terminal MRGS$(H)_6$ and a C-terminal FLAG tag via unique *Bam*HI *x Hind*III sites containing *lacIq* for expression control. After transformation into *E. coli* XL1-blue, 380 single DARPin clones were expressed in 96-well format and lysed by addition of B-PER Direct detergent supplemented with 0.4 mg/ml lysozyme and 20 U/ml nuclease (Pierce). These bacterial crude extracts of single DARPin clones were subsequently used in a Homogeneous Time Resolved Fluorescence (HTRF)-based screen to identify potential binders. Binding of the FLAG-tagged DARPins to the streptavidin-immobilized biotinylated C2a domain of AC9 was measured using FRET (donor: Streptavidin-Tb cryptate (610SATLB, Cisbio), acceptor: MAb Anti FLAG M2-d2 (61FG2DLB, Cisbio)). FRET signals were recorded using a Varioskan LUX Multimode Microplate (Thermo Scientific). From the identified binders, 32 clones were sequenced and 21 single clones identified. To investigate binding to the full-length AC9 an additional ELISA with immobilized AC9 and AC9-Gαs was performed using bacterial crude extracts.

The DARPins were expressed in small scale, lyzed with Cell-Lytic B (Sigma) and purified using a 96-well IMAC plate (HisPur™ Cobalt plates, Thermo Scientific). After IMAC purification, DARPins were analyzed at a concentration of 10 µM on a Superdex 75 5/150 GL column (GE Healthcare) using an Akta Micro system (GE Healthcare) with PBS containing 400 mM NaCl as the running buffer.

#### Protein expression and purification

***AC9.*** The methods for bovine adenylyl cyclase 9 (AC9, Uniprot ID E1BM79_BOVINE, with a C-terminal YFP-twinStrep tag) expression and purification were similar to those previously described (*5*). Briefly, AC9 was expressed using the BacMam system (*6*). HEK293F cells were grown in suspension at 37°C to a density of ~2-2.5×$10^6$ $ml^{-1}$ in Protein Expression Medium (PEM). The cells were collected by centrifugation and resuspended in Freestyle 293 expression medium. The P2 baculovirus for AC9 expression was added to the cells, followed by an incubation for 3-5 hours. The PEM medium was added back to the cell suspension and the cells were further grown for ~60 hours. For AC9 purification, the cell pellets were resuspended in buffer A (50 mM Tris-HCl, pH 8.0, 150 mM NaCl) supplemented with protease inhibitors (1 mM benzamidine, 1 µg/ml leupeptin, 1 µg/ml aprotinin, 1 µg/ml pepstatin, 1 µg/ml trypsin inhibitor and 1 mM PMSF). Cells were lysed using a Dounce homogenizer and the total cell membranes were collected by ultracentrifugation (Ti45 Rotor, 35000 rpm, 1 h). The membranes were solubilized by adding 1 % DDM and 0.02 % cholesteryl hemisuccinate (CHS). After the second round of ultracentrifugation, the supernatant was mixed with 1 ml of CNBr-activated Sepharose coupled with anti-GFP nanobody (*7*), followed by a 30 min incubation with rotation. The resin was collected and washed with 40 column volumes of buffer B (50 mM Tris-HCl, pH 8.0, 150 mM NaCl, 0.1 % digitonin). The protein was eluted by adding HRV 3C protease (1:10 w/w) overnight. The eluted protein was concentrated using a 100 kDa cutoff Amicon concentrator and applied to the Superose 6 Increase 10/300 GL column pre-equilibrated with buffer B. The peak fractions corresponding monomeric AC9 were concentrated and used for cryo-EM grid preparation.

***AC9$_{1250}$.*** The C-terminally truncated bovine AC9$_{1250}$ (residues 1-1250 of AC9) was expressed using transient transfection. HEK 293F cells were grown in suspension at 37°C to a density of ~2-2.5×10$^6$ ml$^{-1}$ in Protein Expression Medium (PEM). The cells were collected by centrifugation and resuspended in Freestyle 293 expression medium. The transfection mixture was prepared containing plasmid and polyethyleneimine (linear PEI max, Polysciences) at a ratio of 1:3 (w/w). After a 3-5 h incubation, PEM medium was added back to the cells and the cells were cultured for ~60 hours. The purification procedure for AC9$_{1250}$ was identical to that for AC9.

***Gαs subunit.*** The procedure for expression and purification of the bovine Gαs (Uniprot ID P04896-1, with a C-terminal 8xHis-tag) was similar to the one previously described (*5*). The standard Bac-to-Bac baculovirus expression system was used (Invitrogen): High Five insect cells were cultured in suspension (1 L) to a cell density of 1.5×10$^6$ ml$^{-1}$ and the P2 virus was added to infect the cells for protein expression. The cells were harvested after 72 hours. For protein purification, the cells were resuspended in buffer A, lysed using a Dounce homogenizer and solubilized by adding 1% dodecyl-β-maltoside (DDM) for 1 hour. The lysate was clarified by ultracentrifugation. The supernatant was incubated with 1 ml Ni-NTA resin for 30 min. The resin was washed with buffer C (50 mM Tris-HCl, pH 8.0, 150 mM NaCl, 0.02 % DDM) supplemented with 20 mM imidazole. This was followed by a second wash with buffer C supplemented with 40 mM imidazole. The protein was eluted with buffer C supplemented with 250 mM imidazole. The eluted protein was concentrated using a 10 kDa Amicon concentrator and applied to a Superdex 200 Increase 10/300 GL column pre-equilibrated with buffer B. The peak fractions were concentrated, snap-frozen in liquid nitrogen in aliquots, and stored at -80 °C until the day of the experiment.

***DARPins.*** BL21(DE3) cells were transformed with a plasmid encoding a DARPin of interest with an N-terminal 8xHis-tag and a C-terminal Flag-tag. The cells grown overnight, and 1 L of LB medium was inoculated and grown at 37 °C to an $OD_{600}$ of 0.8. To induce protein expression, 1 mM IPTG was added to the culture and the induced cells were further grown at 37°C for 3 hours. After harvesting by centrifugation, the cells were resuspended in buffer A supplemented with 1 mM PMSF and lysed by sonication. The lysate was clarified by centrifugation and the supernatant was incubated with 1 ml Ni-NTA resin for 30 min. The resin was washed with buffer A supplemented with 20 mM imidazole, followed by a second wash with buffer A supplemented with 40 mM imidazole. The protein was eluted using buffer A supplemented with 250 mM imidazole, applied to a Superdex 200 Increase 10/300 GL column pre-equilibrated with buffer A. The peak fractions were concentrated, snap-frozen in liquid nitrogen in aliquots, and stored at -80 °C until the day of experiment.

**AC9-C4 complex.** Upon elution of AC9 from the Sepharose-GFP nanobody resin, the protein was mixed with the purified DARPin C4 at a molar ratio of 1:2. The complex was incubated for 30 min at 4°C and applied to the Superose 6 Increase 10/300 GL column pre-equilibrated with buffer B. The fractions corresponding to the AC9-C4 complex were collected and concentrated for cryo-EM grid preparation.

#### Cryo-EM sample preparation

***AC9-M, AC9-C4-M and AC9-C4-A.*** The solutions containing the purified AC9 or AC9-C4 at a concentration of about 5 mg/ml were incubated with 5 mM $MnCl_2$, 0.5 mM MANT-GTP (0.5mM ATPαS for AC9-C4-A sample) for 30 min at 4°C. A small aliquot of the sample (3.5 μl) was applied to the glow-discharged UltraAuFoil 1.2/1.3 300-mesh grid. The grid was blotted for 3 s, plunge-frozen in the liquid ethane using Vitrobot Mark IV (Thermo Fisher Scientific). The grids were transferred to and stored in liquid nitrogen for subsequent cryo-EM data collection.

***$AC9_{1250}$-Gαs-M.*** The purified of $AC9_{1250}$ at a concentration of 5mg/ml was incubated with 5 mM $MnCl_2$, 0.5 mM MANT-GTP and a 2-fold molar excess of GTPγS-activated Gαs protein for 30 min at 4°C. A small aliquot of the sample (3.5 μl) was applied to the glow-discharged Quantifoil R1.2/1.3 200-mesh grid. The grid was blotted for 3 s, plunge-frozen in the liquid ethane using Vitrobot Mark IV (Thermo Fisher Scientific). The grids were transferred to and stored in liquid nitrogen for subsequent cryo-EM data collection.

#### Cryo-EM data collection and processing

The cryo-EM datasets were collected using a Titan Krios electron microscope equipped with a K2 Summit direct electron detector (Gatan) and a GIF-Quantum energy filter (slit width of 20 eV) at EMBL Heidelberg. The micrographs were recorded using SerialEM (*8*) in super-resolution mode. The defocus range was set from -0.75 μm to -2.5 μm. Each micrograph was dose-fractionated to 40 frames with a total exposure time of 8 s, resulting in a total dose of about 40 $e^-/Å^2$.

The movies were binned two-fold and motion-corrected using MotionCor2 (*9*), yielding micrographs with a pixel size of 0.81 Å. Gctf (*10*) was used for CTF estimation. About 2000 particles were first manually picked and 2D classified in Relion 3.0 (*11*). The selected 2D classes were used as templates to autopick the particles in all selected micrographs (AC9: 8608 micrographs, AC9-C4: 8481 micrographs, $AC9_{1250}$-Gαs-M: 10312 micrographs). The initial model of AC9 was generated from PDB: 2HYD using EMAN2(*12*) and lowpass filter to 40 Å. The refined map of AC9 was used as the initial model for 3D reconstruction of AC9-C4. For the 3D reconstruction of $AC9_{1250}$-Gαs-M, the map of AC9-Gαs (EMD-4719) was used as the initial model. In each case, after several rounds of 2D and 3D classifications the best 3D classes were selected and 3D refinement and post-processing were performed in Relion3.0. Local resolution maps were calculated by ResMap (*13*) implemented in Relion 3.0. The detailed steps of image processing for each dataset are shown in Fig. S1, S5-S6, S8 and Table S1.

The AC9-C4-A dataset was collected using Titan Krios electron microscope equipped with a K3 direct electron detector at ScopeM at ETH Zurich. The micrographs were recorded using EPU in super-resolution mode. The defocus range was from -1 μm to -2 μm. Each micrograph was dose-fractionated to 40 frames with a total exposure time of 0.7s, resulting a total dose of about 48 $e^-/Å^2$. All data processing was performed according to the procedure described above for AC9-C4-M.

#### Model building and refinement

Model building was carried out manually in COOT (*14*). The model of AC9-Gαs (PDBID: 6R3Q) was used as reference. The AC9 domains M1, M2, C1a and C2a were fitted individually according to the cryo-EM map of each dataset. The side chains of the amino acids were adjusted based on the clearly defined densities of the bulky residues (Phe, Trp, Tyr and Arg). MANT-GTP was built using COOT. The model was refined using real_space_refine in PHENIX (*15*). Each model was validated as previously described (*5*). Briefly, the atoms of the final models were randomly displaced by 0.5 Å using PDB tools implemented in PHENIX. The perturbed models were refined in PHENIX against the half map1. The refined models were used to generate FSC of model versus half map2. The map vs model FSC curves agreed well for all structures. The geometries of the models were validated using MolProbity (*16*). All figures were prepared in PyMol (*17*) or Chimera (*18*).

**Molecular dynamics simulation**

The catalytic domains of AC9, $AC9_{C1a}$ and $AC9_{C2a}$, in complex with Gαs (Gαs-C1a-C2a) were used for molecular dynamics (MD) simulations. Two ATP conformations (A1 and A2) were manually built in COOT, based on the density elements corresponding to possible conformations of the bound MANT-GTP (M1 and M2) molecules present in the $AC9_{1250}$-Gαs-M reconstruction. The MD system was prepared using solution builder in CHARMM-GUI (*19*). The model was neutralized by adding ions (with the Monte-Carlo method). The water box type was rectangular with an edge distance of 20 Å. The system was solvated with TIP3P water molecules. The CHARMM36m force field was used for the simulations and Gromacs 2019.2 (*20*) was used as the MD engine. The energy minimization was performed with a Fmax tolerance of 1000 kJ $mol^{-1}$ $nm^{-1}$. The system was equilibrated for 125 ps. The trajectory was accumulated for 50 ns with a timestep of 2 fs. The trajectory data were analysed by VMD (*21*) and visualized using VMD and PyMOL.

**Adenylyl cyclase activity assay**

Assay of purified adenylyl cyclase activity was performed as previously described (*5, 22*). The reaction system contained 50 mM Tris-HCl, pH 7.5, 150 mM NaCl, 0.1 % digitonin, 5 mM $MnCl_2$, 100 µM total ATP (10 nM [$^3$H] ATP, PerkinElmer NET1189001MC) and 0.01 mg/ml purified AC9. For DARPin characterization, the purified DARPin protein was added to the system to a final concentration of 0.66 µM (corresponding to an AC9 to DARPin molar ratio of 1:10). The reaction was initiated by adding ATP. The reactions were performed at 30°C for 10-30 mins and terminated by addition of 20 µl 2.2 M HCl and incubation at 95°C for 4 min. The samples were transferred to disposable columns filled with 1.3 g aluminum oxide. [$^3$H] cAMP was eluted with 4 ml ammonium acetate (0.1 M) and determined using liquid scintillation counting.

The effect of DARPin C4 on human AC5, human AC6, bovine AC8 and *M. tuberculosis* Rv1625c, in comparison to bovine AC9 and $AC9_{1250}$, was assessed by preparing two reaction mixtures per tested AC: in the presence and in the absence of 10 µM DARPin C4. The final concentration of 0.005 mg/ml was used for AC5, AC6 and AC8; 0.01 mg/ml concentration was used for AC9 and $AC9_{1250}$, and 0.001 mg/ml concentration was used for Rv1625c.

For dose-response curves for DARPin C4- and Gαs-mediated activation of AC9, DARPin C4 or Gαs proteins were used in a final volume of 200 µl at the following concentrations: 0 µM, 0.033 µM, 0.1 µM, 0.33 µM, 0.5 µM, 1 µM, 3.3 µM and 10 µM.

Membrane assay of AC activity from Sf9 cells expressing individual AC isoforms was performed as described (*23*). AC isoforms were stimulated with 50 nM DARPin and/or 50 nM Gαs in the presence of 5 mM $Mg^{2+}$ and 200 µM ATP. DARPin dose-response curves for AC9 were performed in the presence of 5 mM $Mg^{2+}$ or $Mg^{2+}$ plus $Mn^{2+}$ (0.5 mM) and the indicated concentrations of DARPin.

All data were analyzed using GraphPad Prism (*24*). All enzymatic activity assays were performed at least three times (n = 3) unless indicated otherwise. The values were compared using one-way analysis of variance (ANOVA), followed by Tukey's multiple comparisons test implemented in GraphPad Prism.

#### Cell culture for *in vivo* cAMP accumulation assays and FRET analysis

To study the interactions of DARPin C4 with AC9 by Förster Resonance Energy Transfer (FRET) microscopy and to determine the effects of this reagent on the in vivo AC activity of AC9, $1\times10^6$ cells were seeded per well of a 6-well plate in DMEM medium supplemented with 5% FCS and penicillin/streptomycin. The cells were cultured over-night at 37 °C and 5% $CO_2$ and the next day the cells were co-transfected with the plasmids of interest using PEI, at a DNA to PEI ratio of 1:2 (w/w), with the equal amounts of the AC9-YFP and DARPin-C4-CFP plasmids. In case of *in vivo* cAMP accumulation assay, the negative controls included cells transfected with pcDNA3.1 alone, as well as with the individual plasmids encoding AC9-YFP, DARPin-C4-CFP alone. For FRET analysis, the serotonin transporter SERT, tagged with an N-terminal CFP and a C-terminal YFP, C-SERT-Y (*25*), was used as the membrane FRET control, CFP-YFP fusion as cytoplasmic FRET control and YFP-SERT, YFP-SERT with DARPin C4-CFP, DARPin C4-CFP and AC9-YFP were used as negative controls. Whenever cells were transfected with only one test plasmid, pcDNA3.1 was used in 1:1 ratio [w/w] to maintain uniform DNA amounts across all samples. For the transfection procedure, DNA and PEI dilutions were prepared separately, mixed and incubated for 5-10 min before they were added to the cells. In case of the *in vivo* cAMP accumulation assay, the medium was additionally supplemented with 500 µM isobutyl-methylxanthine (IBMX) and the cells were incubated at 37 °C and 5% $CO_2$ for 48 h.

#### In vivo cAMP accumulation assay

The medium was aspirated from each well, and the cells were washed with 1 ml of PBS. Cells were lysed by addition of 1.5 ml of lysis and detection buffer (Cisbio) for 30 min. Subsequently 16 µl aliquots of the lysate were added to the wells of a 384-well plate (PerkinElmer) in triplicates, together with 4 µl of pre-mixed HTRF antibodies (cAMP Gs dynamic kit, Cisbio) and the plate was sealed and incubated at room temperature for 1 hour. The HTRF was measured at 620 nm and 665 nm using PheraSTAR® FSX plate reader (BMG Labtech) using TR-FRET optical module. The HTRF ratio and cAMP concentration were calculated according to the cAMP Gs dynamic kit protocol.

#### FRET microscopy

The day before fluorescence microscopy, 120,000 cells were seeded into ibidiTreat µ-slide 8-well microscopy chamber (ibidi) and allowed to adhere overnight. Before the experiment, the medium was replaced with FluoroBrite™ DMEM medium supplemented with 5% FCS and penicillin/streptomycin.

The cells were imaged using Nikon Ti Eclipse epifluorescence microscope equipped with an ORCA Flash 4.0 camera (Hamamatsu) using a Nikon CFP Plan Apochromat 60x, NA 1.4 objective at 37 °C and 5% $CO_2$. The “three-filter method” of FRET analysis was performed using a SpectraX light engine, equipped with CFP, YFP and FRET excitation and emission band-pass filters (CFP excitation at 430/24 nm, emission at 470/24 nm; YFP excitation at 500/20 nm, emission at 535/30 nm; FRET excitation at 430/24 nm, emission at 535/30 nm) with 200 ms exposure. The images were processed using Image J plugin PixFRET (version 1.8.0_202). The background (BG) and spectral bleed-through (SBT) parameters were determined for the donor and acceptor separately and were used to generate the calculated FRET and FRET efficiency images. The calculated BG values were subtracted from the final donor and acceptor images. The greyscale FRET efficiency image was used to measure FRET efficiency values at the plasma membrane, which was predefined as the region of interest, except for CFP-YFP construct, which is expressed as a cytosolic protein. A line was drawn randomly through the plasma membrane and pixel intensities along the line were plotted. The pixel intensity values from the plasma membrane were averaged and the average used as final FRET efficiency value per cell. FRET efficiencies were expressed as mean ± S.E.M. and were compared using one-way ANOVA followed by Dunnett’s multiple comparisons test. All statistical analysis was performed using GraphPad Prism 8.0.0.

### Supplementary figures

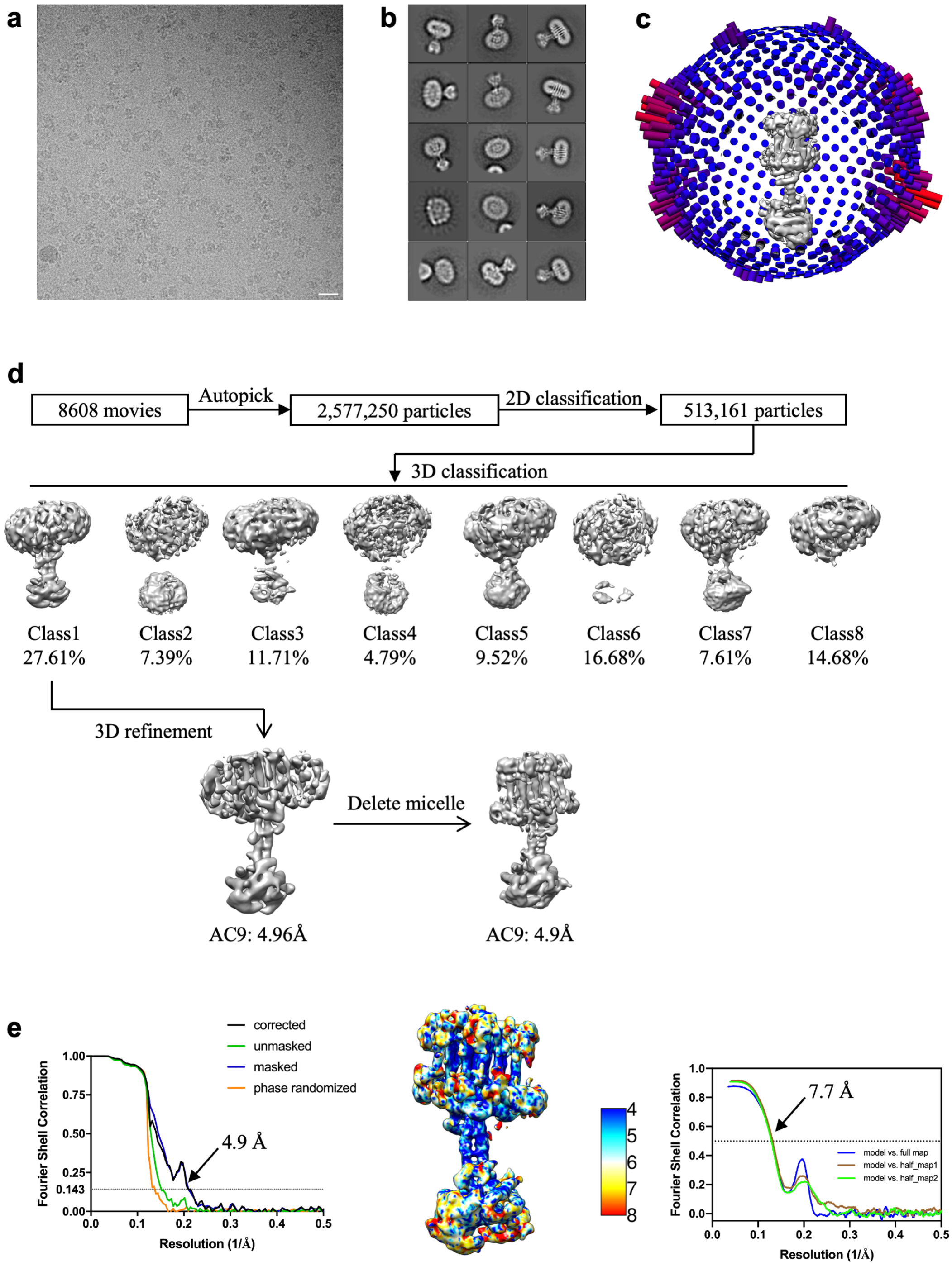


**Supplementary Figure 1 | Cryo-EM image processing procedure of AC9-M dataset. a**, A representative micrograph of AC9 in presence of 0.5 mM MANT-GTP; scale bar corresponds to 20 nm. **b**, 2D classes of AC9-M dataset show distinguishable features, including the detergent micelle and the protruding soluble domain. **c**, Angular distribution of AC9-M dataset. **d**, Overview of the AC9-M dataset image processing procedure, including particle picking, 2D and 3D classification and 3D auto-refinement. **e**, Fourier shell correlation plot (FSC) plots of the map AC9-M (left), local resolution estimation (middle) and map to model FSC plot (right).

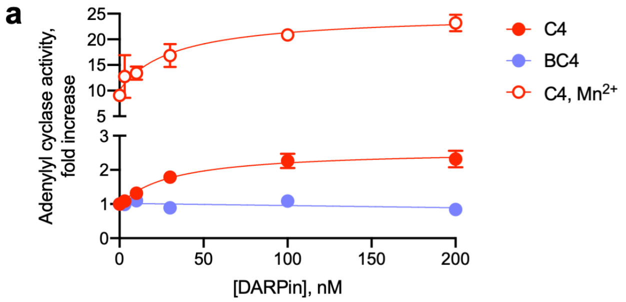


**Supplementary Figure 2 |** Adenylyl cyclase activity assays using AC9 in membrane preparations from Sf9 cells. The DARPin C4 or the negative control, DARPin BC4 (a DARPin selected as a binder for *Mycobacterium intracellulare* Cya/Rv1625c), were titrated into the reaction mixture to determine their effect on AC9 activity. Similar results were obtained in the presence of $Mg^{2+}$ (closed circles) and $Mn^{2+}$ (open circles) as counterions; n = 3 (for BC4 n = 2).

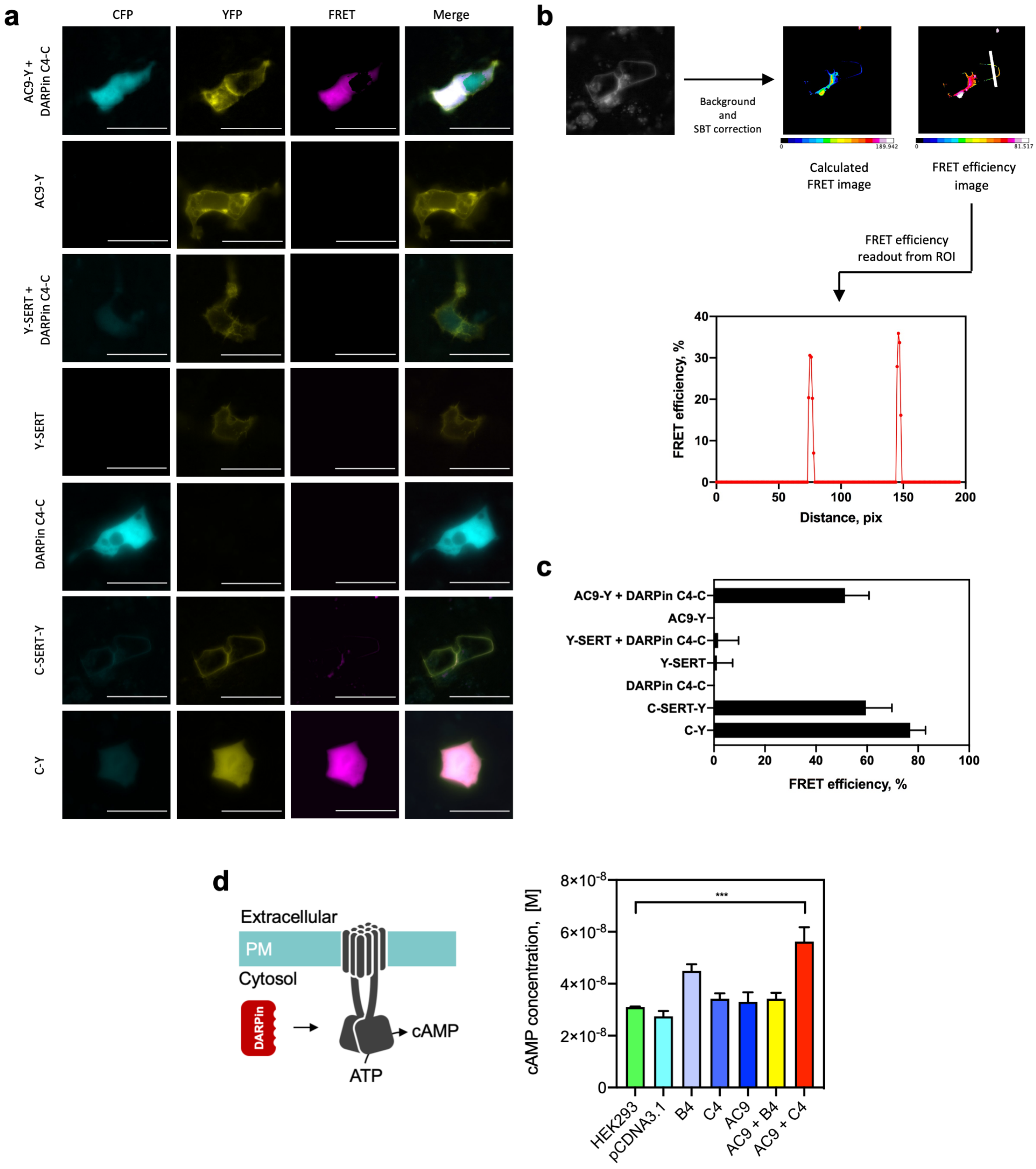


**Supplementary Figure 3 | DARPin C4 interacts with AC9 in vivo. a**, FRET microscopy reveals interactions between AC9-YFP and DARPin C4-CFP in HEK293F cells transfected with the corresponding plasmid DNA constructs. Individual constructs of AC9 and DARPin-C4, along with an unrelated protein, SERT (fluorescently labelled with YFP at the N-terminus), were used as negative controls. CFP-YFP (C-Y) and CFP-SERT-YFP (C-SERT-Y) were used as positive controls. **b-c**, The image processing was performed as described under "Methods"; after background and spectral bleed-through (SBT) correction of the images, the calculated FRET images were used to calculate the FRET efficiency images, regions of interest were defined (corresponding to the plasma membrane for all constructs, and to the cytosolic regions for DARPin-C4-C and C-Y). The maximal FRET efficiency values were averaged and plotted as a bar graph (c). The values in C indicate mean ± S.E.M.; n = 30. **d**, The sketch indicates the activity assays performed using the AC9 and DAPRin-C4 constructs expressed in the HEK293F cells (YFP- and CFP-tagged versions of the proteins, respectively). For all measurements, n = 3 (for B4 n = 2). *Left.* The in vivo cAMP accumulation assays show that only in the presence of overexpressed AC9 and DAPRin C4 together there is significant production of cAMP in the HEK293F cells. One way ANOVA followed by Dunnett's test indicates that out of all tested variations, only the AC9 + C4 sample is significantly different from the control ($P < 0.0005$).

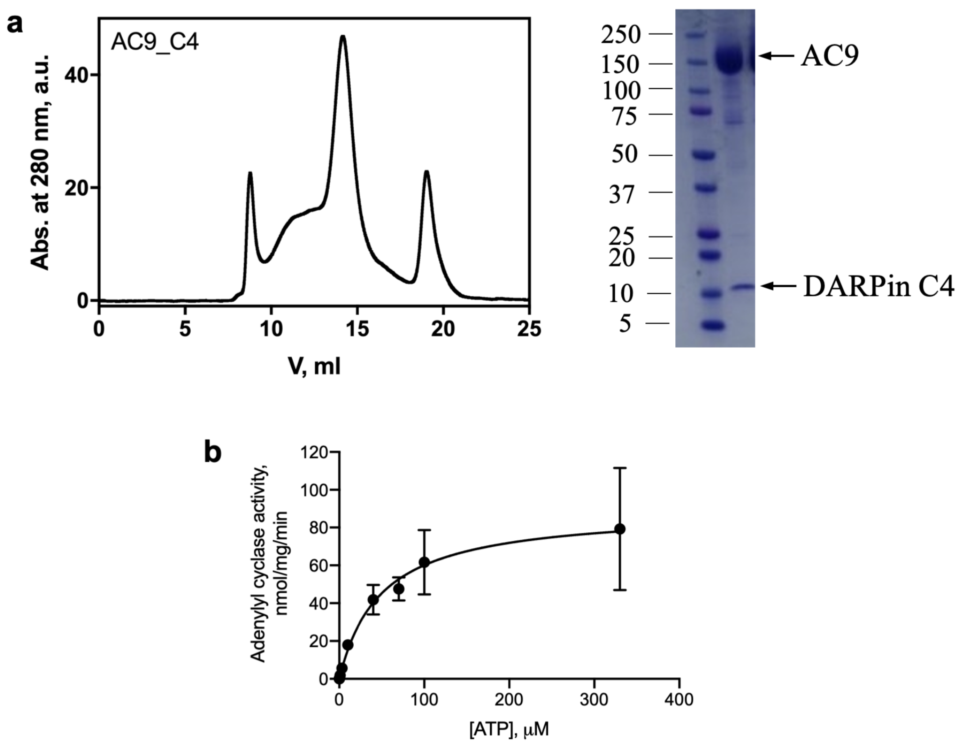


**Supplementary Figure 4 | Purification and enzymatic activity of the AC9-C4 complex. a,** Size-exclusion chromatography (SEC) of the AC9-C4 complex purified in digitonin (*left*) and SDS-PAGE of the purified AC9-C4 complex (*right*). The proteins in the complex are indicated with arrows; the numbers correspond to the molecular weight marker bands on the left side of the gel. **b,** Enzyme kinetics of the AC9-C4 complex reveals a $K_m$ of 49 μM and a $V_{max}$ 90 nmol/mg/min (n = 3).

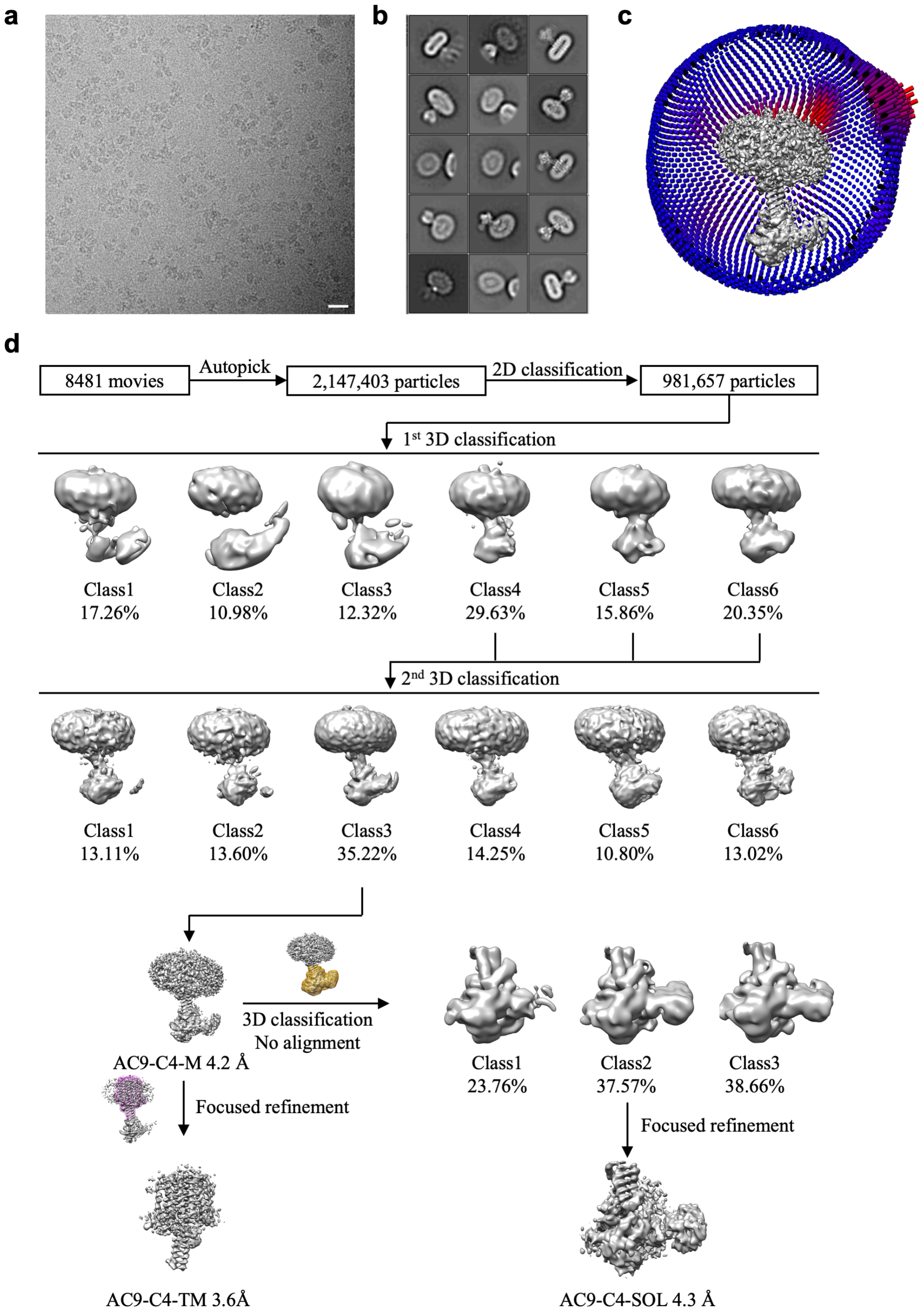


**Supplementary Figure 5 | Cryo-EM image processing procedure for the AC9-C4-M dataset. a**, A representative micrograph of AC9-C4 complex in the presence of 0.5 mM MANT-GTP; scale bar corresponds to 20 nm. **b**, 2D classes of AC9-C4 dataset feature a relatively small density adjacent to the catalytic domain of AC9. **c**, Angular distribution of AC9-C4 dataset. **d**, Overview of the AC9-C4 dataset image processing procedure, including particle picking, 2D and 3D classification, 3D auto-refinement and focused refinement using the mask covering the TM region and the soluble region.

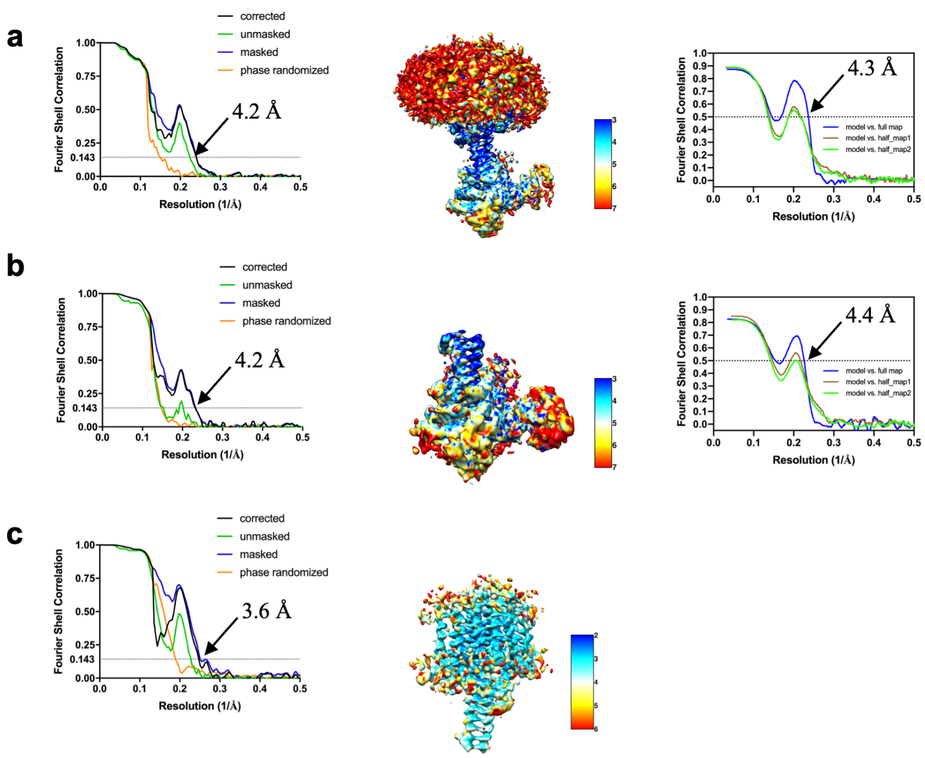


**Supplementary Figure 6 | FSC plots and local resolution estimation for the 3D reconstruction of AC9-C4-M complex.** FSC plots are shown in the left panels, local resolution surface representations for each indicated map estimation are shown in the middle panels, and the map to model FSC plots are indicated in the right panels.

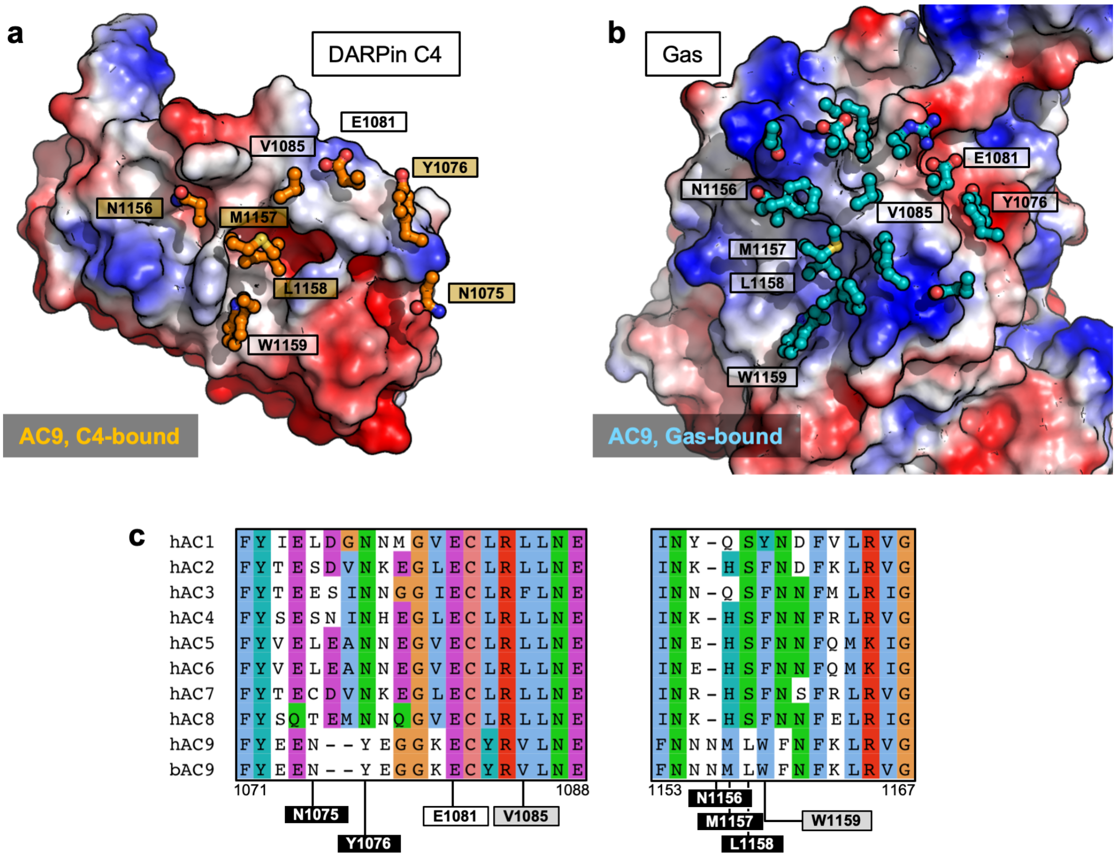


**Supplementary Figure 7 | Unique residues of AC9 at the DARPin C4-binding interface. a**, A view of the DARPin C4, in electrostatic potential-coloured surface representation. The residues of AC9 within 4 Å of the binder are shown as sticks. **b**, Same as in A, with Gαs protein in the AC9-Gαs complex shown in surface representation. **c**, Sequence alignment of the G protein-binding regions in human AC1-AC9 and bovine AC9. The residues unique for AC9 at the DARPin C4-AC9 interface are shown as block boxes (corresponding to the yellow boxes in a). The conserved substitutions are shown as grey boxes.

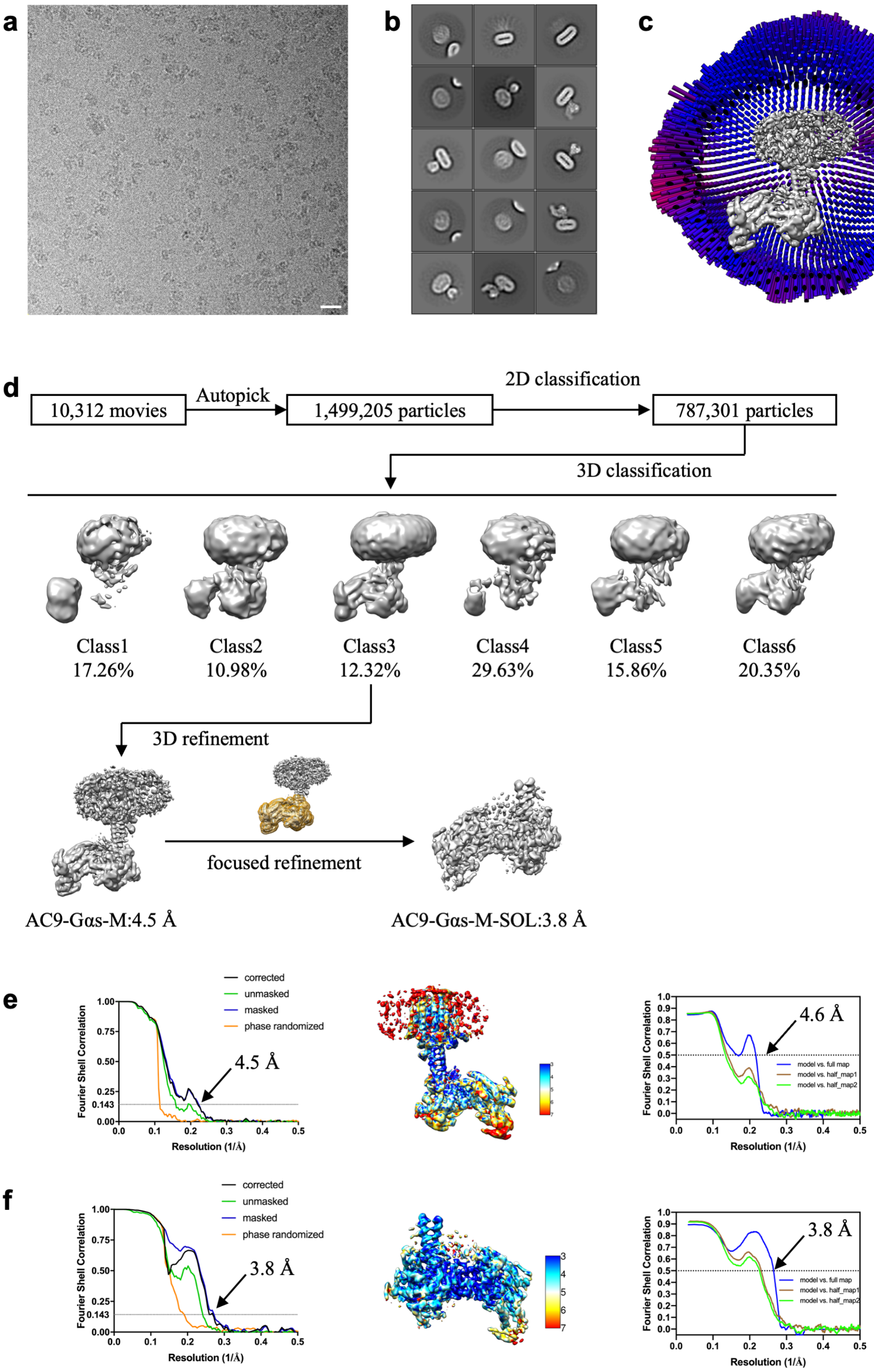


**Supplementary Figure 8 | Cryo-EM image processing procedure of AC9$_{1250}$-Gαs-M dataset. a**, A representative micrograph of AC9$_{1250}$-Gαs complex in presence of 0.5 mM MANT-GTP; scale bar corresponds to 20 nm. **b**, 2D classes of AC9$_{1250}$-Gαs-M dataset. **c**, Angular distribution of AC9$_{1250}$-Gαs-M dataset. **d**, Overview of the image processing procedure, including particle picking, 2D and 3D classification, 3D auto-refinement and focus refinement using the mask covering soluble region of the complex. **e-f**, FSC plots of map (left), local resolution estimation (*middle*) and map to model FSC plots (*right*) of $AC9_{1250}$-G$\alpha$s-M dataset.

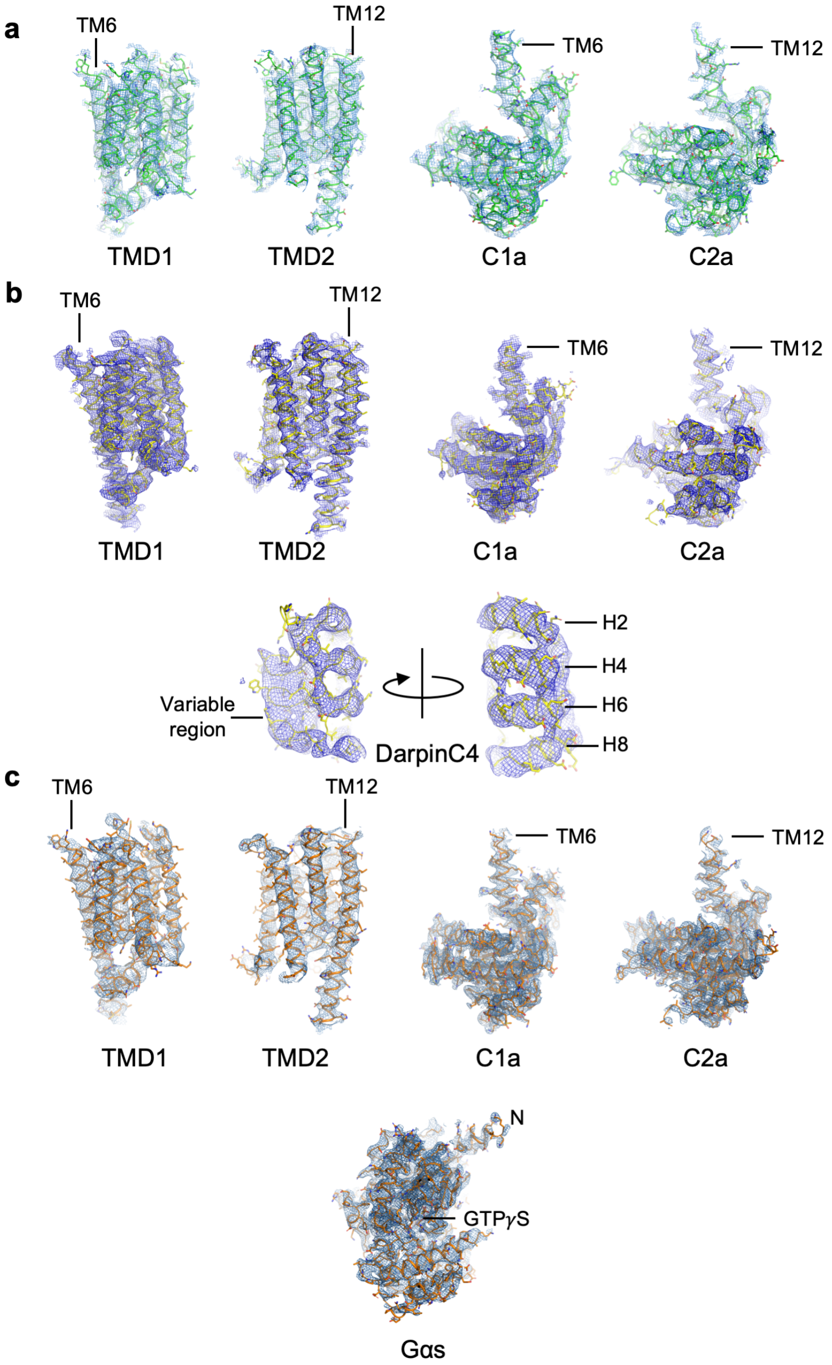


**Supplementary Figure 9 | Features of the cryo-EM density maps.** The density maps in mesh representation and the corresponding models (backbone represented as ribbons and side-chains represented as lines) are shown for: AC9-M (a), AC9-C4-M (b) and AC9-Gαs-M (c). The transmembrane (TMD1, TMD2) and catalytic domains (C1a, C2a), DARPin C4, Gαs subunit are shown seperately. Positions oft he TM6 and TM12 are indicated in the figure.

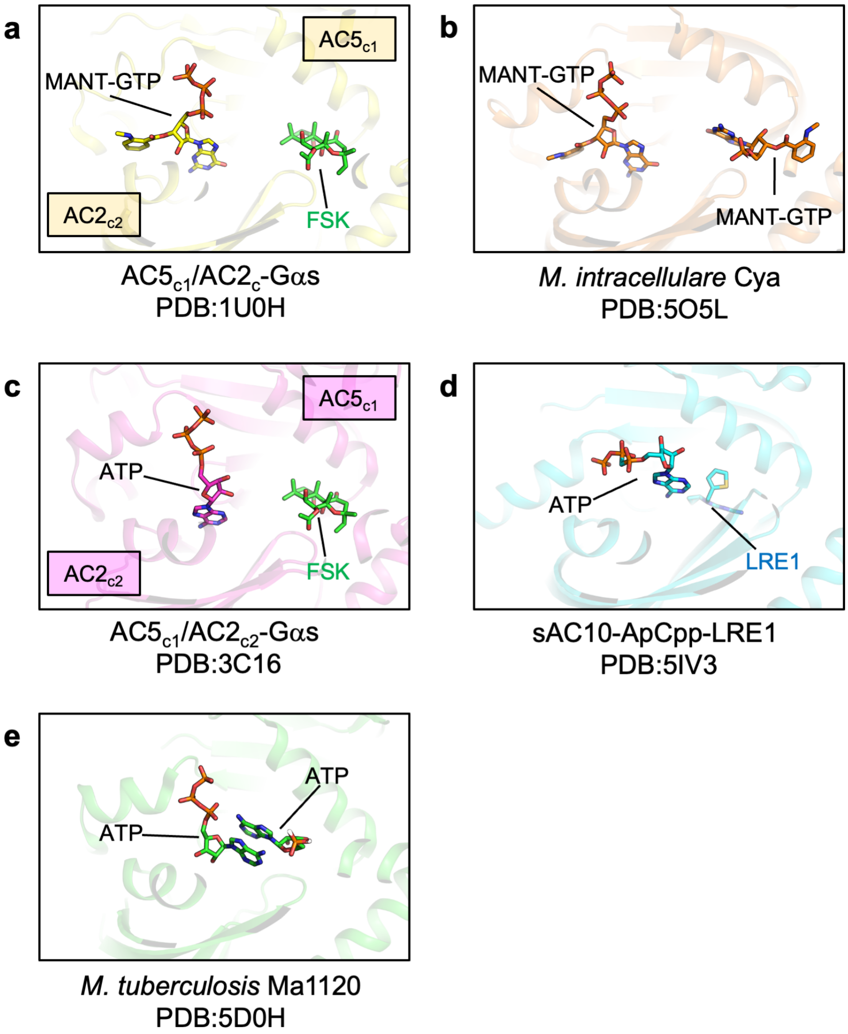


**Supplementary Figure 10 | Comparison of nucleotide-bound states of AC9 with those of other ACs. a**, The crystal structure of $AC5_{c1}/AC2_{c2}$-Gαs (PDB:1U0H) shows MANT-GTP is in M1 pose. **b**, The crystal structure of the *M. intracellulare* Cya (PDB:5O5L) shows MANT-GTP is in M1 pose. **c**, The structure of $AC5_{c1}/AC2_{c2}$-Gαs (PDB:3C16) shows ATP is in catalysis-compatible pose A2. **d**, The crystal structure of sAC-ApCpp-LRE1 (PDB: 5IV3) shows a non-canonical nucleotide pose in which adenine group interacts with the inhibitor LRE1. **e**, The structure of *M. tuberculosis* Ma1120 (PDB: 5D0H) features a non-canonical ATP pose, with adenine moieties of the two adjacent ATP molecules packing against each other.

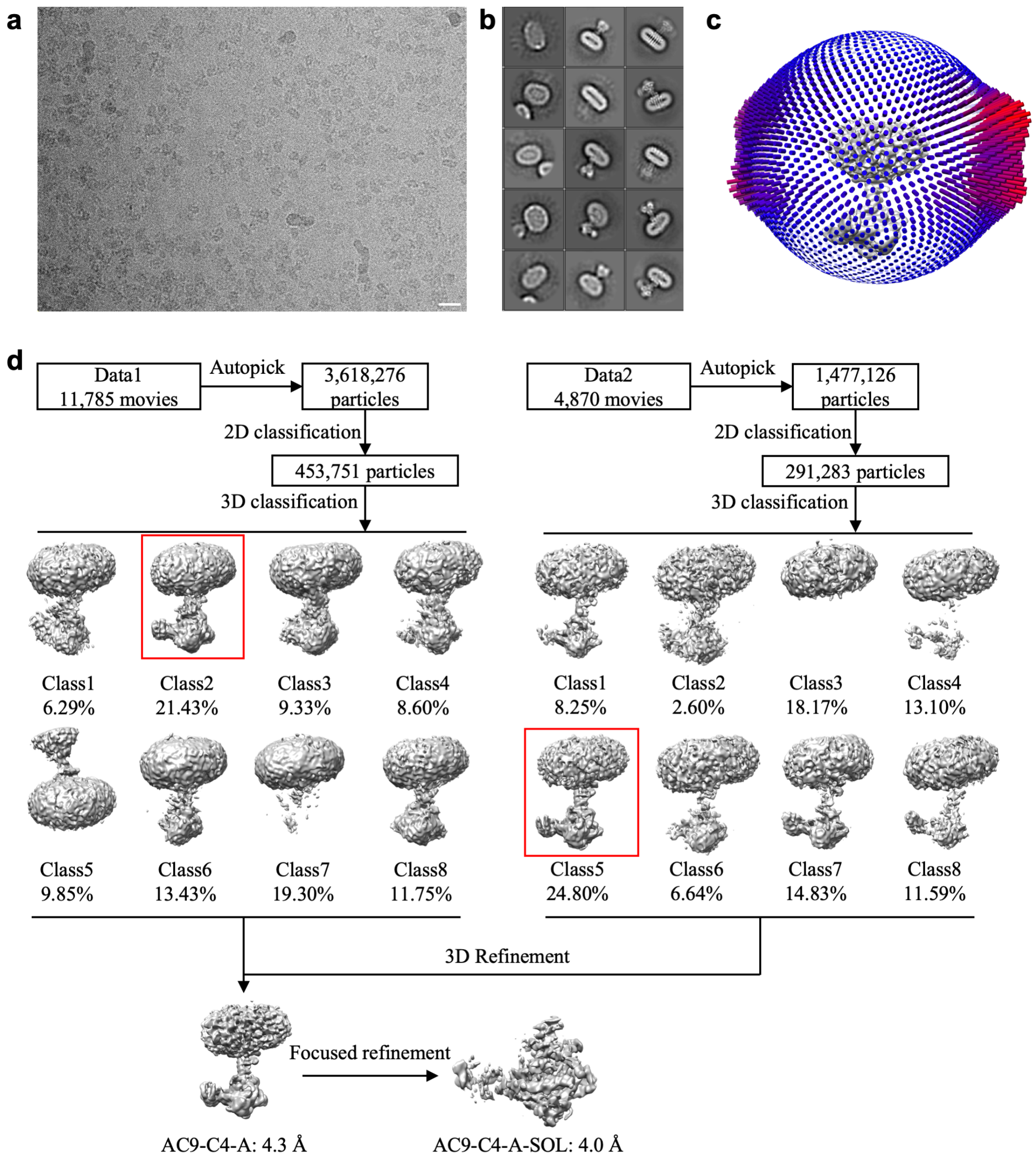


**Supplementary Figure 11 | Cryo-EM image processing procedure for the AC9-C4-A dataset. a,** A representative micrograph of AC9-C4-A complex in the presence of 0.5 mM ATPαS; scale bar corresponds to 20 nm. **b,** 2D classes of AC9-C4-A dataset. **c,** Angular distribution of AC9-C4-A dataset. d, Overview of the AC9-C4-A dataset image processing procedure, including particle picking, 2D and 3D classification, 3D auto-refinement and focused refinement using the mask covering the soluble region.

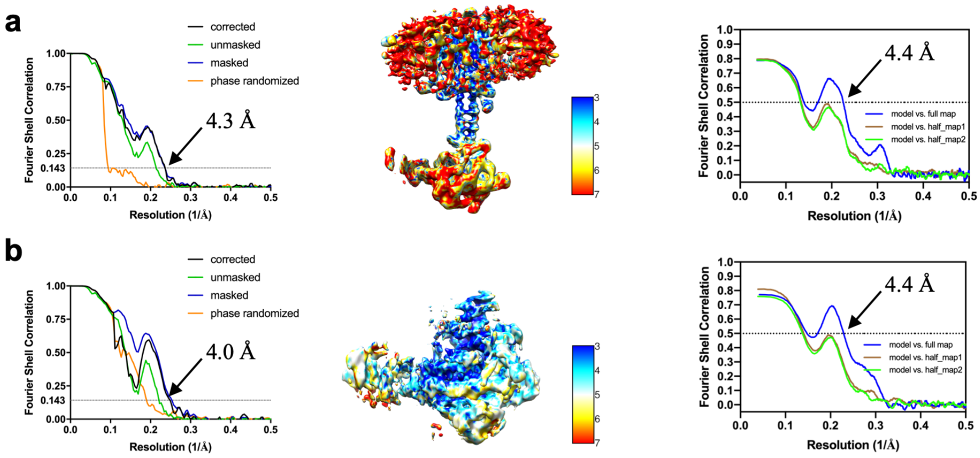


**Supplementary Figure 12 | FSC plots and local resolution estimation for the 3D reconstruction of AC9-C4-A complex.** FSC plots are shown in the left panels, local resolution surface representations for each indicated map estimation are shown in the middle panels, and the map to model FSC plots are indicated in the right panels.

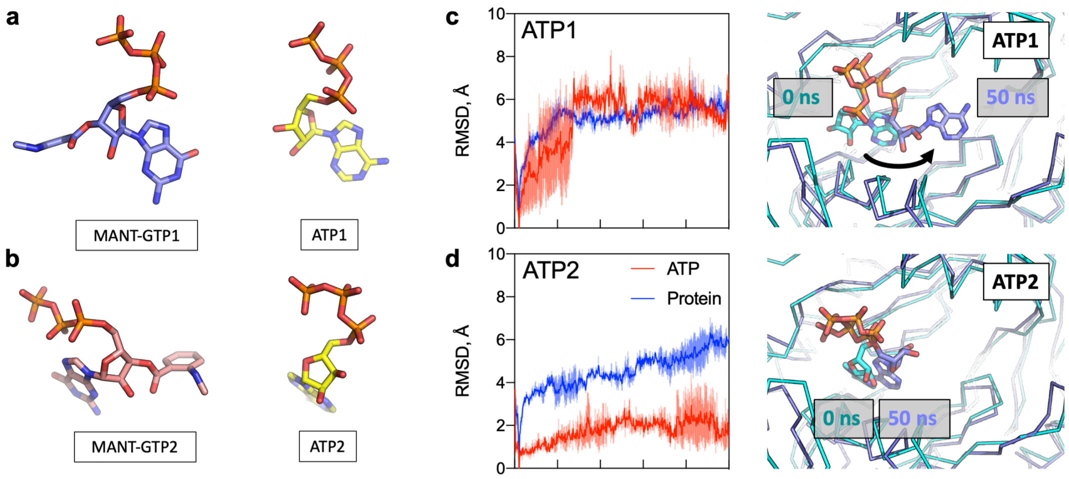


**Supplementary Fig. 13 | Molecular dynamics simulation of the two possible ATP-bound states of AC9. a-b**, The experimentally determined positions of the MANT-GTP molecules correspond to the theoretical substrate-bound (ATP-bound) states of AC9. **c-d**, The models of the AC9 catalytic domains bound to the molecules of ATP in two possible conformations (ATP1 and ATP2), modelled according to the experimentally determined structures of AC9 complexes, were subjected to MD simulations. The traces correspond to the atomic displacements of the protein (blue) or ATP molecules (red). The panels on the right indicate the starting (teal) and the end-states (blue) of each system after 50 ns equilibration. Although state ATP2 (d) shows a relatively stable conformation of ATP, the state ATP1 changes substantially, resulting in long-range displacement of ATP away from the active site (black arrow in c).

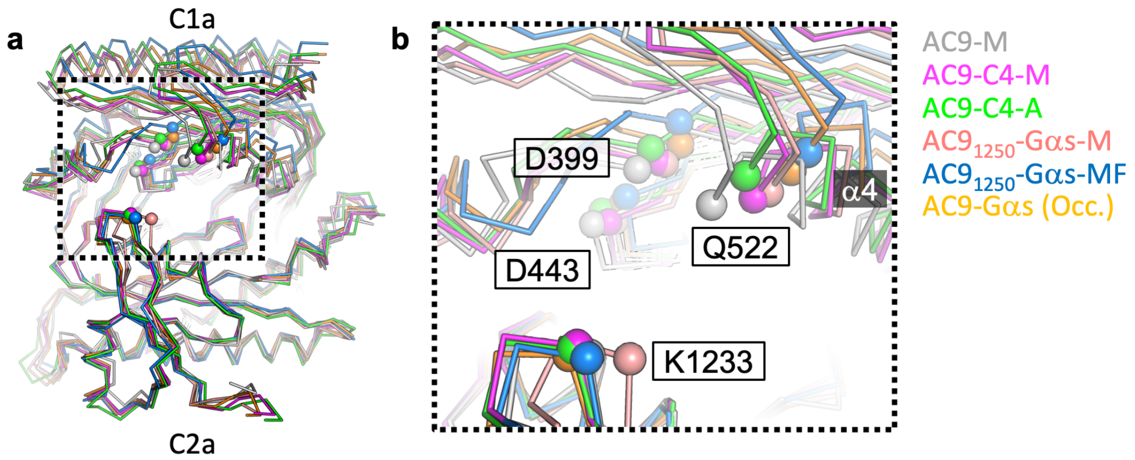


**Supplementary Fig. 14 | Structural transitions in the distinct activator-bound states of AC9 in the presence of MANT-GTP and ATPαS. a**, All available structures of AC9 were structurally aligned using the C2a domain as an anchor (as in Fig. 4e, with an additional structure, AC9-C4-A, coloured green). **b**, Comparison of the five available structures reveals relative displacement of the active site residues, D399, D443, K1233 and the residue Q522 in the helix α4 (Cα atoms are shown as spheres). The individual structures are indicated on the right side of the panel. Comparison of the MANT-GTP- (AC9-C4-M, magenta) and ATPαS-bound state (AC9-C4-A, green) shows a high degree of similarity between them. The distance between the Cα atoms of D399, D443 and Q522 residues in the C1a domain of these two structures is 1.1 Å, 1.4 Å and 1.8 Å, respectively.

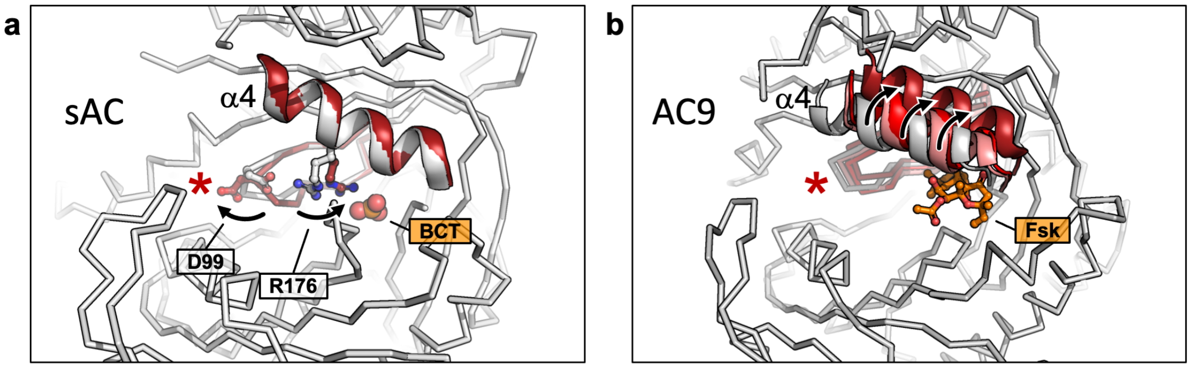


**Supplementary Fig. 15 | Comparison of the activator-induced conformational changes in AC9 and in soluble AC, sAC. a**. In the case of sAC, the helix α4 undergoes minimal changes upon binding of the bicarbonate ion (red α4 helix - bicarbonate-bound state PDB ID: 4cll; white - apo-state PDB ID: 4clf; the protein model in ribbon representation corresponds to the apo state). Instead of going through large-scale domain movements, a side chain of the residue R176 is drawn away from the side chain of D99, a residue that participates in coordinating the metal ions critical for catalysis. This switch mechanism underlies sAC activation by $HCO3^-$. **b**, The conformational changes in AC9 involve whole-domain rearrangements, with helix α4 (which participates in forskolin binding) moving concomitantly with the potency of the activating agent. The ribbon model corresponds to the AC9-M state. The progressively darker red α4 helices correspond to the states AC9-C4-M, AC9-Gas-M and AC9-Gαs-MF. The asterisk in a and b indicates protein activation.

**Supplementary Table 1 | Cryo-EM data collection and refinement statistics.**

| **Data collection** | | | | | | | |
|---|---|---|---|---|---|---|---|
| Sample | AC9-M | AC9-C4-M | | AC9$_{1250}$-Gαs-M | | AC9-C4-A | |
| Instrument | FEI Titan Krios/Gatan K2 Summit/Quantum GIF | | | | | Gatan K3 | |
| Voltage | 300 | | | | | 300 | |
| Electron dose(e$^-$/Å) | 40 | | | | | 48 | |
| Defocus range (μm) | -0.75 to -2.5 | | | | | -1.0 to -2.0 | |
| Pixel size (Å) | 0.81 | | | | | 0.678 | |
| Map resolution (Å) FSC threshold 0.143 | AC9-M | AC9-C4-M-FL | AC9-C4-SOL | AC9-Gαs-M-FL | AC9-Gαs-M-SOL | AC9-C4-A-FL | AC9-C4-A-SOL |
| | 4.9 | 4.2 | 4.2 | 4.5 | 3.8 | 4.3 | 4.0 |
| Number of particles | 141,446 | 210,729 | 210,729 | 157,200 | 157,200 | 170,731` | 170,731 |
| Refinement | | | | | | | |
| Model resolution (Å) FSC threshold 0.5 | 7.7 | 4.3 | 4.4 | 4.6 | 3.8 | 4.4 | 4.4 |
| Map sharpening b-factor (Å) | -192.2 | -204.7 | -189.0 | -197.83 | -183.8 | -100 | -164.169 |
| Map CC | 0.73 | 0.78 | 0.75 | 0.8 | 0.82 | 0.66 | 0.69 |
| Model composition | | | | | | | |
| Protein residues/ligand | 842 | 970 | 577 | 1187/1 | 794/1 | 970 | 577 |
| ADP (B factor) | 351.53 | 149.98 | 138.11 | 205.14 | 76.56 | 199.29 | 164.22 |
| Bond length r.m.s.d. (Å) | 0.005 | 0.009 | 0.004 | 0.006 | 0.008 | 0.005 | 0.002 |
| Bond length r.m.s.d. (°) | 0.866 | 1.075 | 0.741 | 0.779 | 0.818 | 0.819 | 0.644 |
| Validation | | | | | | | |
| MolProbity score | 2.66 | 2.62 | 2.26 | 2.48 | 2.27 | 2.07 | 1.93 |
| Clash score | 38.14 | 33.32 | 20.58 | 29 | 16.33 | 16.04 | 11.3 |
| Rotamer outliers (%) | 0 | 1.09 | 0 | 0.19 | 0.29 | 0 | 0 |
| Ramachandran plot | | | | | | | |
| Favored (%) | 88.29 | 88.78 | 92.95 | 90.76 | 89.9 | 94.76 | 94.71 |
| Allowed (%) | 11.71 | 11.22 | 7.05 | 9.24 | 10.1 | 5.24 | 5.29 |
| Disallowed (%) | 0 | 0 | 0 | 0 | 0 | 0 | 0 |

**Movie 1 | Molecular dynamics simulation of ATP1 state.** ATP1 shifts from its original location.

**Movie 2 | Molecular dynamics simulation of ATP2 state.** ATP2 shows a relatively stable conformation in 50 ns of the simulation.

**Movie 3 | Molecular morph of AC9 in different states.** AC9-C4 (magenta, partial activated state), AC9-G$\alpha$s (orange, occluded state), AC9$_{1250}$-G$\alpha$s-M (pink, partial activated state) and AC9$_{1250}$-G$\alpha$s-MF (blue, fully activated state) are aligned to AC9-M (white, basal state) based on C2a domain. The whole-domain rearrangements correspond to increased levels of AC9 activation.
